# Recreating the Biological Steps of Viral Infection on a Bioelectronic Platform to Profile Viral Variants of Concern

**DOI:** 10.1101/2023.11.11.566634

**Authors:** Zhongmou Chao, Ekaterina Selivanovitch, Konstantinos Kallitsis, Zixuan Lu, Ambika Pachaury, Róisín Owens, Susan Daniel

## Abstract

Viral mutation rates frequently outpace the development of technologies used to detect and identify harmful variants; for SARS Coronavirus-2 (SARS-CoV-2), these are called variants of concern (VOC). Given the continual emergence of VOC, there is a critical need to develop platforms that can identify the presence of a virus and readily identify its propensity for infection. We present an electronic biomembrane sensing platform that recreates the multifaceted and sequential biological cues that give rise to distinct SARS-CoV-2 virus host cell entry pathways and reports the progression of entry steps of these pathways as electrical signals. Within these electrical signals, two necessary entry processes mediated by the viral Spike protein, virus binding and membrane fusion, can be distinguished. Remarkably, we find that closely related VOC exhibit distinct fusion signatures that correlate with trends reported in cell-based infectivity assays, allowing us to report quantitative differences in fusion characteristics among them that inform their infectivity potentials. This cell-free biomimetic infection platform also has a virus-free option that equally reports infectivity potential of the Spike proteins. We used SARS-CoV-2 as our prototype, but we anticipate that this platform will extend to other enveloped viruses and cell lines to quantifiably explore virus/host interactions. This advance should aid in faster determination of entry characteristics and fusogenicities of future VOC, necessary for rapid response.

## Introduction

RNA viruses tend to have high mutation rates than their DNA counterparts^1–3^, hence developing vaccines and antivirals that remain protective against disease-causing viral variants remains challenging with the fast pace of emerging variants of concern (VOC). When outbreaks of new viruses occur, quickly establishing the mechanisms of viral entry and transmission are critical for the rapid development of vaccines and therapeutics to combat RNA viruses and assessing emerging VOC and determining their potential for human harm. However, doing this is no easy feat; the mechanisms of viral infection are complex, involving numerous, multi-step biological processes, which often vary across cell types and microenvironments, hence necessitating a protracted incubation period for comprehensive data acquisition by live cell-based assays^4–6^. The entry pathway chosen by SARS-CoV-2, for example, is highly dependent on the interactions at the host cell membrane-virion interface, as well as the local protease, pH, and ionic conditions^7^. Because viral entry represents the first contact viruses have with host cells, the proteins and mechanisms that comprise these events have been targeted therapeutically and diagnostically to block or detect viral infection.

The last few years have witnessed a surge of advancements aimed towards the rapid, sensitive, and accurate detection of viruses and their emerging variants. While Reverse Transcription Polymerase Chain Reaction (RT-PCR)^8^ remains the gold standard for detection, other classical methods include antibody detection^9^, which relies on detecting antigen specific antibodies in serum, and antigen detection^10^, which uses designer antibodies to bind to and detect antigens. While these methods have been instrumental throughout the COVID-19 pandemic, they provide a binary response indicating either a detectable presence or absence of an antigen but offer few insights into their infectivity potential and are not appropriate for screening VOC. Furthermore, studies have shown that as variants emerge, the ability of designed primers and antibodies to maintain their sensitivity diminishes, requiring the detection materials to be reformulated^11^. CRISPR-Cas– and isothermal amplification-based detection technologies have also been developed^12,13^. Techniques that fall into both categories detect nucleic acid (RNA or DNA) sequences and, while offering high sensitivity and selectivity, do not offer insights into a virus’ structural integrity. Biosensors, on the other hand, have been shown to differentiate between a virus protein and a whole virus particle. They have been successfully used as detection platforms for coronaviruses by exploiting the specificity of antigens for their respective receptors^14–17^. However, to comprehensively understand the unique properties of emerging mutants and their potential for infection beyond mere binding interactions, a functional assessment of infectivity potential is imperative.

For enveloped viruses, which contain a lipid membrane that wraps or “envelopes” the genome-filled capsid, infection of the host cell is initiated when virions first bind to a host cell receptor, followed by the triggered fusion of the viral membrane with that of the host. These critical entry steps (binding and fusion) allow for the viral genome to be delivered to the host’s cytosol. Chemically-responsive glycoproteins that protrude from the viral envelope mediate these entry processes. Their interactions with the cell plasma membrane receptors and other chemical cues create a fusion-promoting microenvironment. The cues that lead to viral entry typically involve some sequence of exposure to receptors, specific proteases, low pH, and ions. Depending on the host cell type, the identity of the triggers and the sequence of their presentation can vary. Additionally, the glycoprotein’s properties (*i.e.,* mutations that alter the glycoprotein in some way) also influence how they respond to these cues. Thus, it is a complicated convolution of glycoprotein sequence and host cell environment that control the entry of these viruses and ultimately create conditions for a productive infection of the host. Coronaviruses (including SARS-CoV, MERS and SARS-CoV-2), are a family of enveloped viruses and the variety of conditions that influence their biological entry pathways represent a major hurdle in probing viral infection mechanisms *in vivo*, as many methods lack the necessary control of the microenvironmental conditions and clear signals of a successful entry process. To gain the upper hand in mitigating future virus outbreaks and staying ahead of emerging VOC, it is crucial to develop platforms that are capable of both mimicking infection conditions and reporting infection progress via quantifiably readouts.

Here, we demonstrate the power of a new technique that can detect viral entry processes, but importantly, provide quantitative readouts that distinguish entry characteristics of closely-related viral strains. Starting with SARS-CoV-2 Wuhan-Hu-1 (WH1) as a model target, we present the design of an *in vitro*, cell-free entry platform (with a virus-free option as well) based on supported lipid bilayers (SLBs) that faithfully replicates the conditions that promote entry but in a convenient, controllable, and tailorable format with a much faster response time than live cell assays. This cell-free platform senses entry functions electrically and is thus label-free. Next, we probe the entry characteristics of two SARS-CoV-2 Omicron subvariants, Omicron BA.1 and Omicron BA.4, and show that our platform identifies the known differences in fusion activity between these strains as well as repeats the known trends in their cell infectivity. This demonstration of using bioelectronics for detecting and characterizing virus-host entry processes is a critical precursor of the events that lead to host infection. Our device, mimicking the earliest events in “*infection-on-a-chip”,* opens possibilities for examining entry conditions that can be leveraged for both basic science studies, screening assays for antiviral therapies, and fast assessment of entry characteristics that can inform next steps in combatting VOC as they emerge.

## Results

### A description of the biological pathways of SARS-CoV-2 entry that are recreated in this platform

The exterior glycoprotein of SARS-CoV-2 is called Spike^18^. After the initial binding event between Spike and the host cell receptor (membrane-bound angiotensin-converting enzyme 2 (ACE2), viral entry continues via one of two entry pathways depending on its spatiotemporal exposure to microenvironmental cues^19^. Fig. 1 briefly summarizes the two identified pathways for SARS-CoV-2 infection and the critical extra– and intracellular conditions that distinguish them, specifically focusing on the initial steps of binding to, and fusion with, the host cell’s membrane. Which one of two entry pathways is triggered is cell type specific and based on the availability of proteases for S2 cleavage. The first pathway, referred to here as the early entry pathway, is initiated when the transmembrane serine protease 2 (TMPRSS2) is present in the plasma membrane of the host cell^20^. Upon the binding of Spike protein to ACE2, the Spike is cleaved by TMPRSS2 to initiate virus-host membrane fusion presumably at or near the cell surface and the viral genome is delivered to the cytosol. The second pathway, referred to here as the late entry pathway, proceeds when the membrane-bound protease is not present in the plasma membrane of the host cell^21^. In this scenario, Spike protein binds to ACE2 and the virion is endocytosed, where it is subsequently cleaved by the endosomal cysteine protease-cathepsin L (CatL) inside the low pH microenvironment of the endosome. These cues trigger the fusion of the viral envelope with the endosomal membrane and release the genome into the cytosol.

**Fig. 1.**
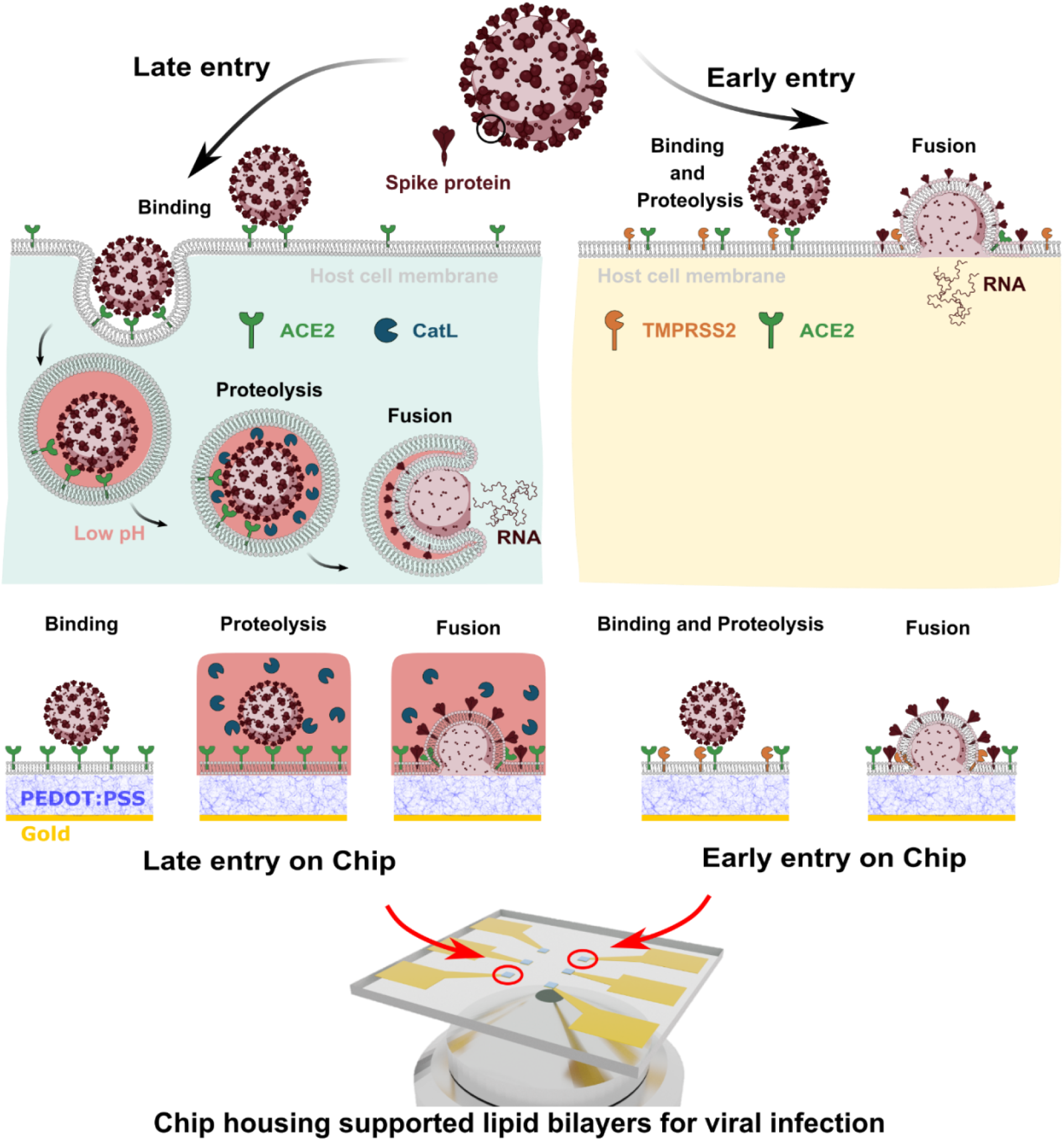
SARS-CoV2 entry pathways and the components needed to recapitulate these entry routes in an *in-vitro* platform. The two known pathways of SARS-CoV-2 including early entry, in which fusion is triggered by the TMPRSS2 protease, and late entry, in which virus particle fusion is catalyzed by the protease CatL at low pH (note: pink color = acidic environment). We propose SLBs self-assembled on PEDOT:PSS electrode provide an ideal *infection-on-chip* platform. The SLB is formed using cell-derived blebs and fusogenic vesicles on PEDOT:PSS surface, hence the membrane components are preserved. Viral pseudoparticles (VPP) with Spike protein, pH swap and soluble catalyst (CatL) can be included to induce fusion. The optically transparent and conductive nature of PEDOT:PSS also allowed both optical and electrical readouts to identify trends characteristic of binding and fusion events.

### Design Parameters for an *Infection-on-Chip* Device

Taking inspiration from biological mechanisms, we set out to design a platform that can faithfully reproduce the microenvironments needed to selectively trigger either of the two entry pathways, and thus initiate the primary steps in a viral infection. There are four essential components in the construction of this *infection-on-chip* platform: 1) the presentation of viral and host cell membrane components, 2) spatiotemporal control over environmental cues required for triggering fusion, 3) a biocompatible scaffold accommodating membrane components for successful infection and 4) quantifiable readouts reflective of different infection stages.

To test the *infection-on-chip* platform for its ability to recapitulate specific cell membrane environments that induce CoV entry events, we used Spike_WH1_-incorporated viral pseudoparticles (VPP_WH1_), produced using previously established methods^22^. Confirmation of Spike protein incorporation is included in Supplementary Fig. 1. To capture the host cell features required for entry, but in a cell-free format, we selected SLB to serve as host cell membrane mimics. These SLBs were composed from native cell membrane components (collected as plasma membrane blebs, or cell blebs) and “fusogenic” lipid vesicles, which self-assemble into a planar, single-bilayer lipid membrane blended with native cell membrane components, *i.e.,* ACE2 receptors and TMPRSS2 proteases. The fusogenic vesicles used in this work are reconstituted from purified 1-oleoyl-2-palmitoyl-*sn*-glycero-3-phosphocholine (POPC) lipids, which is the principal lipid component of mammalian and viral membranes. As shown in Fig. 1, this versatile, easy-to-assemble biomimetic membrane allowed spatiotemporal control over environmental cues to recapitulate both early and late entry pathways: when cell blebs containing ACE2 and TMPRSS2 (confirmed as shown in Supplementary Fig. 1) are incorporated into the SLB, colocalizing receptors and membrane proteases, the early entry pathway can be accessed. When SLBs are formed with only ACE2 containing blebs (confirmed as shown in Supplementary Fig. 1), only fusion via late entry pathway can be activated when CatL is added under acidic conditions.

SLBs can be readily self-assembled on various functional supports, including biocompatible conductive polymers, which prompted our design of a label-free *SLB-on-electrode* structure to directly translate the interactions occurring at the biomimetic membrane into an electrical readout. Our group has previously demonstrated that SLBs can form on PEDOT:PSS (poly(3,4-ethylenedioxythiophene) polystyrene sulfonate) supports^23,24^, a conductive, transparent polymer mixture widely used in biosensing applications^25,26^. We have also demonstrated that SLBs prepared on these polymer supports maintain two-dimensional fluidity of the constituents (both lipids and membrane proteins) and that the buffer-swollen polymer serves as a cushion that supports this characteristic of cell membranes in the resultant SLB^24^.

Presented in the following sections, we fulfill all design parameters necessary for capturing SARS-CoV-2 *infection-on-chip* in a cell-free and label-free manner by building a biomembrane bioelectronic platform. We demonstrate that the label-free electrical readouts of this platform can provide a quantitative approach that could be used for investigating emerging variants, identifying potential variants of concern, and potentially thwarting the progression of outbreaks. For example, the platform could be used as a tool to discover means to disrupt or arrest viral entry processes in anti-viral drug development, or in efforts to classify and differentiate properties of emerging variants as strains evolve, which can assist in predicting host tropism susceptibilities, or inform next generation formulations of vaccines.

### Characterization of the *Infection-on-Chip* Device

The sizes of VPP_WH1_, cell blebs, and synthetic POPC vesicles were measured using Dynamic Light Scattering (DLS) and Nano Particle Analysis (NTA), and are reported to be approximately 100-200 nm, 150-450 nm, and 100 nm, respectively (Supplementary Fig. 2 and Supplementary Fig. 3). The particles count, provided by NTA analysis, allowed us to control the relative concentrations of the VPP_WH1_ and blebs used to assemble the SLBs.

The method of forming SLBs from cell blebs and POPC vesicles on a PEDOT:PSS support is described in **Methods**. To assess their formation, we used fluorescence recovery after photobleaching (FRAP) measurements to confirm the formation of a mobile bilayer on PEDOT:PSS-coated glass coverslips — a critical prerequisite for the fusion events described later in this paper. For this optical characterization, SLBs formed on PEDOT:PSS surface were labeled with a lipophilic dye, R18, and in the case of a mobile bilayer, the R18 dye should diffuse freely throughout the SLB plane. Fig. 2a shows typical FRAP data showing the full recovery of a photobleached spot on a SLB assembled using Vero E6 cell blebs and POPC vesicles. Vero E6 cells were chosen due to their endogenous expression of ACE2; therefore, the SLBs formed using blebs derived from this cell line incorporated the ACE2 receptor^27,28^. The other SLBs assembled for our study were derived from recombinant Vero E6 cells containing the TMPRSS2 receptor, and HEK 293 cells used to assemble Spike-incorporating SLBs. We chose a TMPRSS2-modified Vero E6 cell line for the early entry pathway to remain consistent across as many parameters as possible for comparison with the late entry pathway using Vero E6 cells^29,30^. The FRAP images for these SLBs on PEDOT:PSS can be found in Supplementary Fig. 4 and Supplementary Fig. 5. Upon photobleaching, the fluorescent intensity as a function of time in the photobleached spot was collected and fit with a Bessel function expression (see **Methods**) to later calculate the diffusion coefficient, D. All SLBs show comparable diffusion coefficients: ranging from 0.16-0.2 μm^2^/s and 0.92-0.99 mobile fractions (Supplementary Fig. 6).

**Fig. 2.**
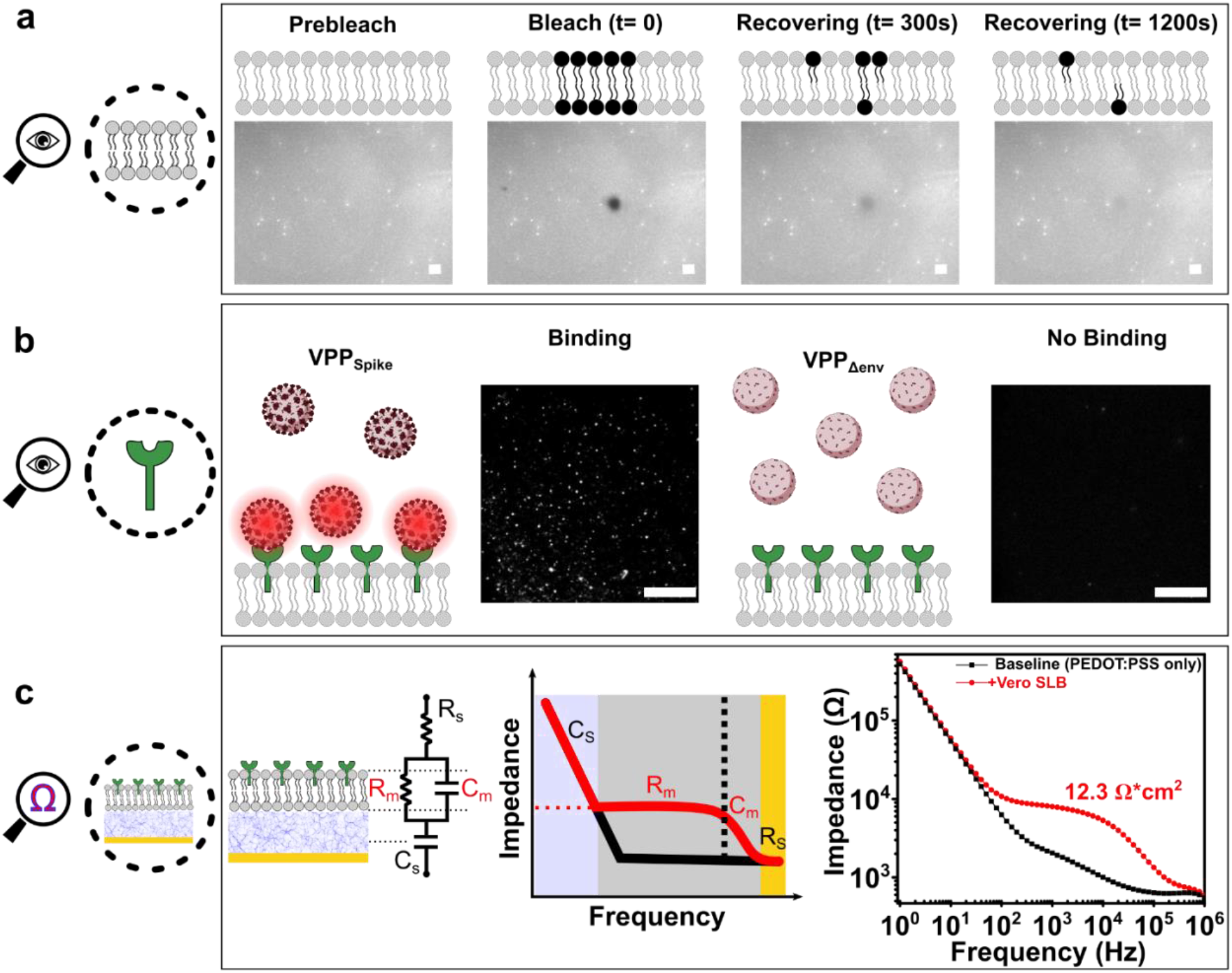
Optical and electrical characterization of the *infection-on-chip* platform’s components and functionalities. (**a**) We used FRAP to characterize the mobility of SLBs formed on PEDOT:PSS surfaces. Shown here is a photobleached spot recovering over time, indicating a mobile SLB. The cartoon representation is meant to provide a conceptual illustration of the technique. Indeed, our SLB was composed of both fluorescent and non-fluorescent lipids and the fluorescence seen in the images are reflective of only the doped in R18 dye. (**b**) TIRF was used to confirm the existence of ACE2 receptor in SLBs: only fluorescently labeled VPP_Spike_ are visible at the SLB interface when bound to ACE2 receptors, while no fluorescently labeled VPP_Δenv_ were observed near the SLB due to the lack of binding interaction with ACE2 receptors on SLB. (**c**) EIS was used to characterize the electrical properties of an SLB on a PEDOT:PSS electrode. An SLB is modeled electrically as a capacitor and a resistor connectedly in parallel, hence its resistance (R_m_) can be extracted by fitting into the RC(RC) circuit as shown, it can then be normalized by the area of electrode. All scale bars in this figure represent 20 μm.

To confirm the ACE2 receptors were incorporated into the SLBs, additional characterization of our SLBs was conducted using total internal reflection fluorescence (TIRF) microscopy. TIRF is an optical imaging technique especially suited to study the interactions occurring near the SLB-bulk interface, as its induced evanescent wave illuminates a limited (∼ 100 nm) vertical region from this interface, effectively eliminating fluorescence from the bulk emanating from unbound virus particles. Although our ultimate goal here is to validate a label-free sensing platform for virus, the visualization of binding events between ACE2 and VPP_WH1_ was necessary to verify that native cell receptors from blebs were incorporated into the SLB assembled on PEDOT:PSS. VPP_WH1_ were labeled with R18 fluorophores that partition into the VPP membrane envelope; the SLB was not labeled in this experiment. We used TIFR microscopy to measure specific interaction between ACE2 receptors in the SLBs and fluorescently labeled VPP_WH1_. As shown in Fig. 2b, a representative TIRF field of view (FOV) provided evidence that the R18-labeled VPP_WH1_ are specifically bound to the ACE2 assembled SLB, while particles devoid of Spike proteins (VPP_Δenv_) do not exhibit any detectable signals, as a control case.

PEDOT:PSS is not only conductive, it is also a volumetric capacitor — making it an ideal electrode material for electrochemical impedance spectroscopy (EIS) measurements as it significantly reduces system impedance^31^. Lower system impedance enables the measurement of small changes in SLB electrical properties that can be correlated with viral entry processes, as we describe later. EIS is a non-invasive electrical sensing technique with a proven track record for accurately quantifying bio-recognition events occurring at biointerfaces^32–34^. When a SLB is self-assembled on PEDOT:PSS electrodes, the ionic flux reaching the electrode surface is reduced due to SLB shielding, thereby decreasing the ionic current. This outcome is ultimately measured by an increase in circuit impedance when compared to the electrode baseline signal (a circuit without SLB coating). The PEDOT:PSS electrodes used in this work were defined on a gold contact pad using optical lithography (see **Methods**). As shown in Fig. 2c, when no SLB was formed on the PEDOT:PSS electrode, the circuital response to alternating voltage with changing frequency is plotted in black, generating a “hockey stick” shape bare electrode baseline signal (PEDOT:PSS only), indicating a typical resistor-capacitor in series structure. Upon the self-assembly of a SLB on the electrode (+Vero SLB), the circuital response shifted from black “hockey stick” to red “chair shape”, confirming the addition of a resistor-capacitor in parallel structure — an established electrical trait of lipid bilayers^35,36^. The membrane resistance (R_m_) and capacitance (C_m_) of the SLB can then be extracted by fitting the signal into an equivalent electrical circuit, as depicted, and then normalized by the area of the electrode.

### Recreating the SARS-CoV-2 Entry Pathways using *Infection-on-Chip*

Now that we have formed a mobile SLB with confirmed native membrane components using FRAP, demonstrated that the initial steps in SARS-CoV-2 infection process (*i.e.,* binding) using TIRF, and verified that SLB-formation results in a measurable signal using EIS, we continued to investigate if fusion can be initiated and detected on our chip when environmental cues are integrated spatiotemporally.

We first focused on the early entry pathway where we mimicked the respective fusion triggering environment by forming a SLB from cell blebs containing ACE2 and TMPRSS2 on the PEDOT:PSS surface. We then introduced the VPP_WH1_ to monitor binding and fusion as depicted in the schematic shown in Fig. 3a. Electrical readouts were conducted on PEDOT:PSS electrodes. As expected, when SLBs were formed on the electrodes, the electrical circuital response to alternating voltage shifted from the black (PEDOT:PSS only) to the red (SLB) signal, as shown in Fig. 3a (right). Subsequently, upon the addition of the VPP_WH1_ to the SLB with both ACE2 receptors and TMPRSS2 proteases, the circuital response shifted from red to blue and, when fitted and normalized, the SLB membrane resistance increased from 13.1 to 19.9 Ω*cm^2^ (+ 51.9 %). We hypothesize that this increase in resistance is attributed to the incorporation of additional biomacromolecules originally in the viral envelope now present in the SLB after the fusion event takes place **—** an observation that is consistent with optical data under the same conditions (Supplementary Fig. 7). VPP_WH1_ were also added to a SLB containing ACE2 (no TMPRSS2 protease) to measure the electrical response arising from binding interactions, while VPP_Δenv_ were added to the SLB with both ACE2 TMPRSS2 to identify any non-specific interactions between the bilayer and pseudo particles not directed via Spike-ACE2 binding. The electrical responses from both control groups are consistent with optical data as shown in Supplementary Fig. 7, suggesting minimal interactions when compared to fusion. The differences in both electrical and optical signals between binding and fusion events suggest that we can differentiate between them under conditions suitable for the early entry pathway using the electronic label-free approach on our *infection-on-chip* devices.

**Fig. 3.**
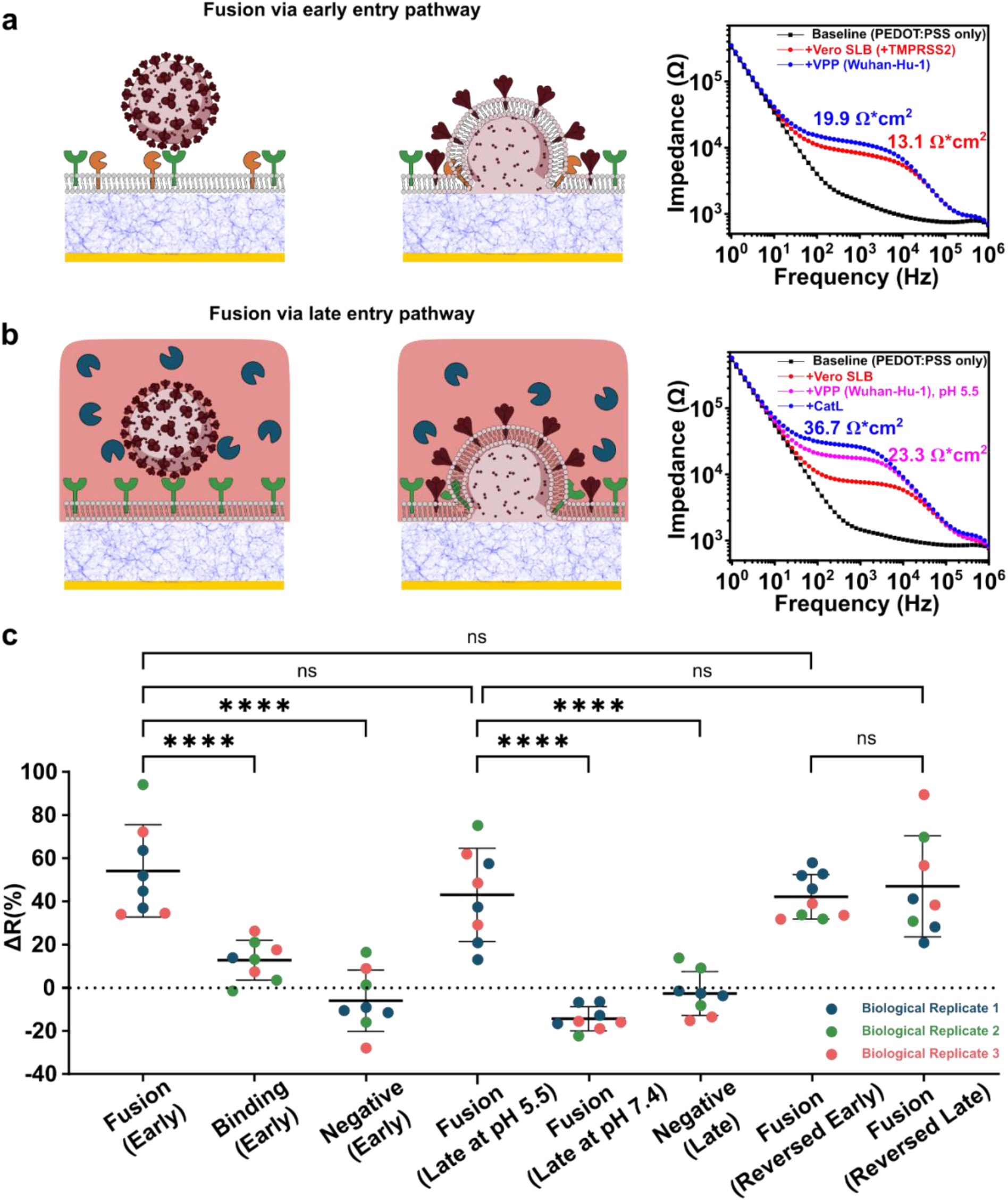
Electrical responses of fusion via early and late entry pathways recapitulated on host cell derived SLB. (**a**) The experimental group for early fusion pathway consisted of VPP_Spike_ and an SLB containing ACE2 (green) and TMPRSS2 (yellow). EIS Signals are characteristic of fusion events showing the changes in SLB membrane resistance in the equivalent electrical circuit scenario; (**b**) the experimental group consisted of the VPP_Spike_ and an SLB containing ACE2 (green) and CatL (navy), where signals are characteristic impedance data of fusion events (note: pink color = acidic environment); **(c)** distribution of membrane resistance changes at all events of all systems (3 biological replicates and n ≥ 8 for all systems) using Spike protein from SARS-CoV-2 Wuhan-Hu-1. ΔR data are mean ± SD; statistical analysis was performed using one-way analysis of variance (ANOVA) with Šidák’s multiple comparisons test, **** (p < 0.0001), ns = non-significant.

The late entry pathway requires protease CatL, instead of TMPRSS2, to catalyze the virus-membrane fusion. We were able to reproduce this pathway using our model system by supplementing the bulk solution with CatL, which is a soluble protein, and mimicking the acidic endosomal environment in which CatL is active. To mimic this environment and triggering conditions in our platform, we generated SLBs made from Vero E6-derived blebs, which contained the ACE2 receptor but no TMPRSS2. To recreate the endosomal triggering environment, as shown in Fig. 3b, we exchanged the initial pH 7.4 buffer to a more acidic buffer (pH 5.5) and then added soluble CatL, which is active at pH 5.5 but not at pH 7.4. Similar to the early entry pathway, the electrode baselines were acquired before SLB formation (black) and after SLB formation (red), as shown in Fig. 3b (right). VPP_WH1_ were first added to bind with the ACE2 receptors in the SLB at pH 7.4, before exchanging the buffer to a more acidic environment (pH 5.5). As a result, the electrical signal shifted from red to pink, indicating that the binding between ACE2 receptors and the VPP, together with the pH drop contributed to an increase in membrane resistance, aligned with our observation in the early entry pathway (Supplementary Fig. 7) and previous report^33^. Upon the addition of CatL, the SLB membrane resistance further increased (blue trace, from 23.3 to 36.7 Ω*cm^2^, + 57.5 %), suggesting successful fusion between the VPPs and SLB membranes. As a control for fusion at non-optimal triggering conditions, CatL was also added to a non-acidic buffer environment after the VPP_WH1_ addition. The electrical signal (Supplementary Fig. 8) suggested membrane resistance dropped insignificantly, indicating there was no fusion due to non-optimal triggering conditions; VPP_Δenv_ were used as a negative control and no significant membrane resistance shift was observed at lower pH after the addition of CatL (Supplementary Fig. 8). These measurements are all congruent with the optical data (Supplementary Fig. 8). From the electrical and optical data it is clear that both CatL and acidic conditions are required for promoting fusion of the VPP_WH1_ with the SLB, an observation that is consistent with our current understanding of SARS-CoV-2 viral entry^37,38^.

The repeatability over biological and technical replicates of electrical responses for fusion and control groups for both pathways is shown in Fig 3c. The change in resistance values for fusion events of VPP_WH1_ are comparable in the early (+ 54.0 ±20.0 %) and late entry (+ 42.9 ±20.2 %) pathways, both distinct from all control groups.

### Differentiating between Wuhan-Hu-1, Omicron BA.1, and BA.4 strains using *Infection-on-Chip*

The VPP_WH1_ were used in all the entry experiments so far and we have confirmed both entry pathways can be recapitulated using the *Infection-on-Chip* platform. Next, we investigated if our platform was capable of distinguishing SARS-CoV-2 variants with different fusogenicities. Omicron BA.1 and BA.4 (BA.1 and BA.4) were selected in this study since BA.1 has been reported to be less fusogenic than BA.4, while both Omicron variants selected have lower fusogenicities than the WH1^39–41^.

The electrical readouts modeling the early and late entries of BA.1 are shown in Fig. 4a. Interestingly, we see no significant membrane resistance increase in the case of early pathway (left), yet a small, but distinguishable resistance increase can be observed in the case of late pathway (middle). Statistical data (right) suggested significance between early and late entries of BA.1, matching more recent reports^40,42–45^.

**Fig. 4.**
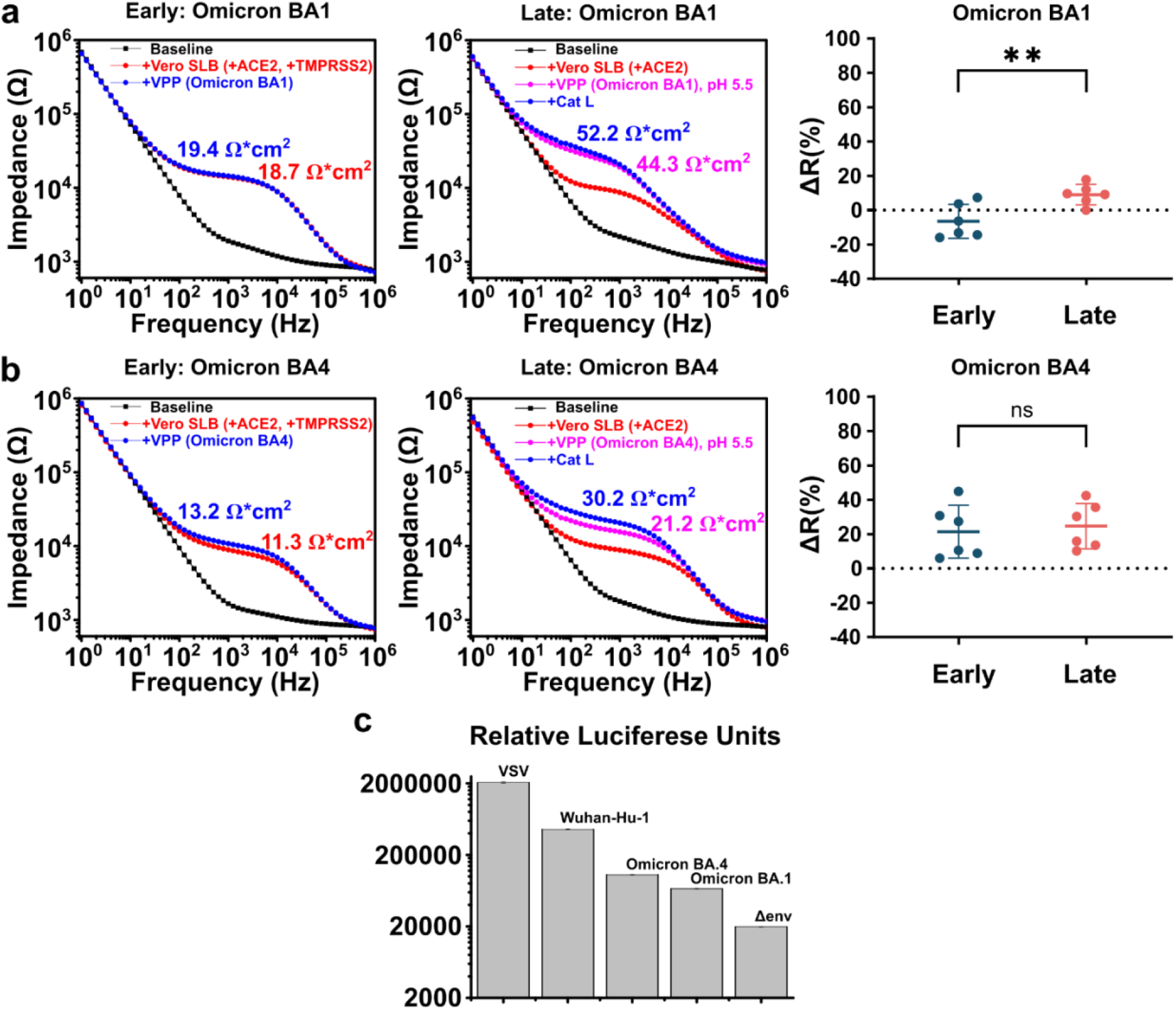
A comparison of viral fusogenicities of SARS-CoV-2 VOC by electrical response and viral transduction assays. **(a)** EIS electrical signal change of fusion via early (left), late (middle) entry pathways and their statistical comparison (right, n = 6 for both pathways) using Omicron BA.1 VPP; **(b)** EIS electrical signal change of fusion via early (left), late (middle) entry pathways and their statistical comparison (right, n = 6 for both pathways) using Omicron BA.4 VPP. ΔR data are mean ± SD; statistical analysis was performed using one-way analysis of variance (ANOVA) with Šidák’s multiple comparisons test, ** (p < 0.01), ns = non-significant; **(c)** relative transduction efficiencies of the Wuhan-Hu-1 Spike and Omicron variant Spike-containing pseudoparticles. The transduction efficiency of VPP_Spike_ was assessed against a positive control that contained a vesicular stomatitis virus G protein (VPP_VSV_) and a negative control without any envelope protein (VPP_Δenv_). The luciferase production of the infectious VPP_Spike_ and VPP_VSV_ was consistently orders of magnitude higher than VPP_Δenv_, indicating that the particles we produced were “active” and capable of fusion with a cell membrane. The samples labeled BA.1 and BA.4 refer to Omicron variants. All infectivity assays were completed with Vero E6 TMPRSS2 cell lines. All data above represent five technical replicates (n = 5). Error bars represent standard deviation.

The resistance values of both early (left) and late (right) entries of BA.4 are shown in Fig. 4b. Comparing BA.1 to BA.4 VPP, membrane resistance increases were more significant for VPP_BA.4_, as suggested by statistical data (right), supporting the reports of BA.4 being more fusogenic than BA.1^39,46^. However, when comparing wild type SARS-CoV-2 to the BA.4 strain, shown in Fig 3c, the membrane resistance increase caused by the fusion of VPP_BA.4_ was still significantly reduced: from + 54.0±20.0 % to + 21.4±10.3 % for the early entry pathway and from + 42.9±20.2 % to + 24.6±12.1 % for the late pathway. Our results matched strongly with viral transduction assays as shown in Fig. 4c, where the relative luciferase units detected using the VPP_WH1_ were about 7x higher than VPP_BA.1_ and 4x higher than VPP_BA.4_. A detailed description of the transduction assay is provided in the **Methods** section. Our EIS based fusion assay aligns well with other standard assays used to determine relative infectivity of virus particles, such as syncytia and plaque assays, evaluating the relative fusogenicities of the three variants explored here^39–41^, confirming the accuracy of *Infection-on-Chip* platform in distinguishing SARS-CoV-2 variants.

### Reversing SARS-CoV-2 Early and Late Pathway Configurations

The previous arrangements used the SLB as a model for either the cellular or endosomal membrane surfaces and the VPP as mimics of the infectious virus. Here, we swap the active constituents of both entry pathways, where now the SLB displayed features found on the virus surface (*i.e.,* the glycoproteins), while blebs in the bulk phase presented their respective host cell surfaces. Specifically, we constructed SLBs that contained Spike_WH1_ protein and formed cell blebs that contained ACE2 or ACE2/TMPRSS2 as host cell “particles” that can bind to and fuse with the Spike-containing SLBs. By swapping the constituent presentation, we present an intriguing strategy for rapidly screening cell types and their respective susceptibility to viral infection without the need for virus particles (virus-free) and only the spike protein gene for cellular expression.

Spike proteins were incorporated into the SLB by rupturing Spike-transfected HEK293 blebs, while TMPRSS2-modified Vero E6 cell blebs (Fig. 5a) and Vero E6 cell blebs (Fig. 5b) were introduced to evaluate and quantify their interactions with the “virus-like” SLB. Mirroring our previous experiments, the SLBs were formed on both PEDOT:PSS coated glass coverslips (Supplementary Fig. 9) and PEDOT:PSS electrodes for optical and electrical readout, respectively.

**Fig. 5.**
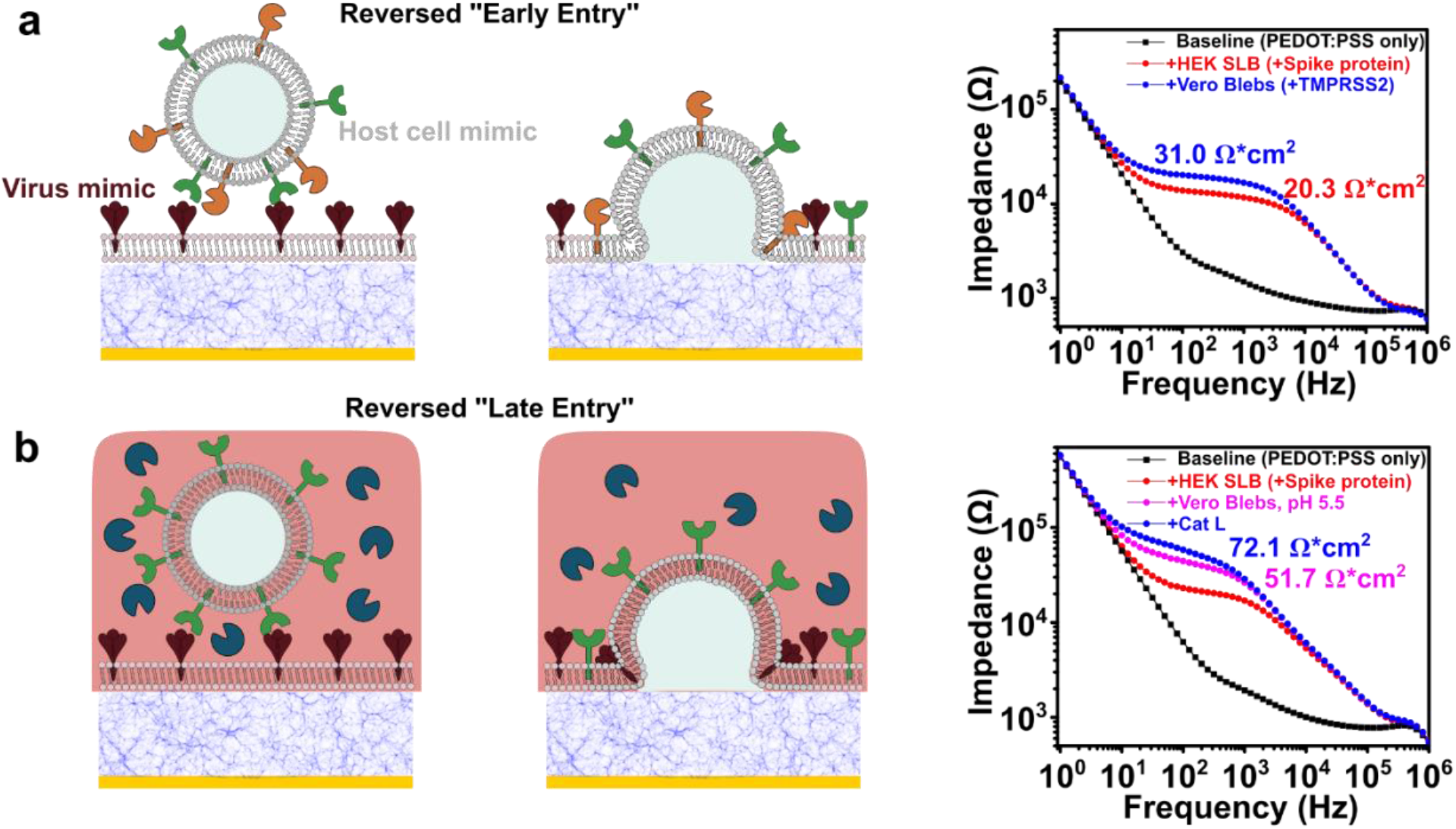
Reverse geometry fusion studies using SLB containing Spike protein with different blebs. (**a**) Illustration of “reversed” early entry pathway and its corresponding EIS electrical signal change; (**b**) “reversed” late pathway and the corresponding EIS electrical signal change.

We first investigated the electrical responses for reversed “early entry”. Similar to the more traditional display of constituents described earlier, the electrical signal shifted from the electrode baseline (black) to the SLB signal (red) (Fig. 5a). After the addition of cell blebs with ACE2 receptors and TMPRSS2, Spike SLB membrane resistance increased from 20.3 to 31.0 Ω*cm^2^ (+ 52.7 %), showing a similar membrane resistance increase as measured in early entry pathway as shown in Fig. 3a and 3c. Similarly, in the reversed “late entry”, cell blebs with ACE2 were added to bind with the Spike SLB and soluble CatL was added to initiate the fusion, after swapping to an acidic buffer environment. The electrical response at each step was measured and plotted in Fig. 5b. After the formation of Spike SLB (red), membrane resistance increased to 51.7 Ω*cm^2^ (pink) upon binding with ACE2-containing blebs and exchanging to a lower pH buffer environment (from PBS pH 7.4 to PBS pH 5.5). Membrane resistance increased to 72.1 Ω*cm^2^ (blue) after the addition of CatL (+ 39.5 %), comparable to the electrical response of the late entry pathway shown in Fig. 3b and 3c. This work shows the SLB based *infection-on-chip* platform can be used to quickly screen interactions between Spike proteins and host cell membranes without producing VPP or virus-like particles (VLP). This can be especially useful for screening antibodies against Spike protein and small molecule fusion inhibitors in a high throughput manner.

## Discussion

### Infection-on-chip Model

The *Infection-on-Chip* devices rely on surface electrodynamics measurements to denote specific interactions between the VPP and the host. To achieve this, the devices were constructed from the necessary biological and chemical elements described earlier and responses were modeled using electrical components of resistors and capacitors. The most rudimentary system resulted in signal contributions from the electrolyte solution resistance and PEDOT:PSS capacitance (electrode baselines in Fig. 2-5). The resistance fluctuations are used as a diagnostic tool to distinguish between the binding and fusion events, the data for which are presented in Fig. 3-5.

Although we have not yet identified the mechanism by which binding and fusion lead to an increase in resistance, we hypothesize that it could be via two potential mechanisms: 1) as more material is integrated into the SLB, the increase in protein and lipid density results in an increase in resistance due to tighter packing, or 2) as more proteinaceous and lipid materials are added to the SLB, membrane defects, or “gaps”, are filled in, consequently increasing the resistance. Both hypotheses are evinced in the fusion pathways for fusogenic WuHan-Hu-1, where resistance values increased by 40-60%. Conversely, the less significant resistance increases upon binding, shown in Fig. 3c and Supplementary Fig. 7, support the second hypothesis. In this scenario, the ACE2 and VPP interactions result in VPP immobilization proximal to the SLB surface, potentially blocking defects near the binding site, without integrating into the SLB itself, though these experiments are still ongoing. Overall, identifying and characterizing fusion events can be especially beneficial for isolating particularly infectious viral variants or screening for therapeutics that target either event.

### SARS-CoV Model System

The two known entry pathways of SARS-CoV-2 capture the canonical features of coronavirus’ initial infection stages, making it an excellent model system for our study. Though the specific receptors and required triggers vary between viral strains and species, there are fundamental aspects that are conserved. For example, there are currently seven identified human coronaviruses (hCoV), among which the most notable are SARS-CoV and MERS-CoV. Though SARS-CoV and SARS-CoV-2 entry mechanisms share more similarities, both requiring an ACE2 protein for binding, all three coronavirus (SARS-CoV-2, SARS-CoV, and MERS-CoV) share similar fusion mechanisms via TMPRSS2 or CatL activation. Going beyond the Coronaviridae family, viruses from the Orthomyxoviridae and Rhabdoviridae families, such as influenza and VSV respectively, also share similarities with the late entry pathways of SARS-CoV-2, requiring an acidification step to prompt fusion. The *Infection-on-Chip* platform may be leveraged to easily identify cell-types particularly susceptible to each virus, provide mechanistic information into the events that initiate infection, and evaluate differences between emerging variants.

The SARS-CoV-2 model system provided an opportunity to determine whether or not the platform can detect variability between different Spike protein variants. Since Omicron variants are now dominant globally, we produced Omicron Spike–incorporating VPP, compared to the Spike_Wuhan-Hu-1_ proteins used for the initial experiments. Experiments were first conducted on Omicron BA.1 variants, which showed a decrease in fusion activity, as evinced by a decreased change in resistance for both early– and late-entry pathways (Fig. 4a), albeit a less significant decrease was observed for the late-entry pathway. Interestingly, this was consistent with recently published findings in which it was identified that Omicron BA.1 and BA.2 variants exhibit an altered entry preference compared to ancestral SARS-CoV-2: preferring endosomal (late) entry pathway as these Omicron variants are less dependent on the TMPRSS2 protease^40,42–45^. Since there are several Omicron variants that have emerged, each with unique sets of mutations, we also evaluated a more fusogenic variant **—** Omicron BA.4. Strikingly, our data correlated well with these reports, as the change in resistance increased for both pathways when using VPP incorporating Spike_Omicron BA.4_ (Fig. 4b). Our data was not only analogous to existing reports of entry-pathway preference, we were also able to detect fusion variability between WH1, Omicron BA.1, and Omicron BA.4 variants that directly mirrored those reported^39^. In these reports the WH1 exhibited the highest fusogencity, followed by Omicron BA.4, and Omicron BA.1 as the least fusogenic of these mutants. These distinctions further highlight the benefits of using this platform with electrical sensing for straightforward screening of viral mutants, and use the acquired data to distinguish highly infectious mutants from those that are less infectious.

### Prospects

“Bioprocesses”-on-chip devices, such as cell-on-a-chip, organ-on-a-chip, and tissue-on-a-chip for instance, represent emergent platforms of interest amongst the biomedical and biomaterials community. Among a myriad of other benefits, their recent successes as *in vitro* micro-scale physiological models can potentially transform fields that focus on therapeutic development and personalized medicine. Our proposed infection-on-chip platform complements these existing technologies by providing mechanistic information at the membrane level without relying on downstream effects or signals. In other words, our readouts directly correlate to events at the membrane-virus interface with exquisite control over the participating components (*i.e.,* receptors, environmental conditions, and presented pathogens), how they are presented, and functionality of the participating constituents (i.e. either binding or fusion events between the SLB and VPP). Whether using the more traditional display, in which the SLB is mimicking the cellular surface, or a presentation where the SLB is emulating the viral surface, the *infection-on-chip* platform can be employed as a quantitative scaffold to interrogate biological pathways or as a tool to rapidly screen interactions with a viral or cellular surfaces, both of which should assist determining societal responses as VOC continue to emerge.

## Methods

### Materials

The 1-palmitoyl-2-oleoyl-sn-glycero-3-phosphocholine (POPC), used for the preparation of fusogenic liposomes, was purchased from Avanti Polar Lipids (700 Industrial Park Dr, Alabaster, AL 35007). Biotechnology grade chloroform was used during the preparation of the POPC liposomes and was purchased from VWR (1050 Satellite Blvd. Suwanee, GA 30024). Whatman Nucleopore polycarbonate filters (50 nm) (Cytiva-Marlborough, MA) were used for liposome extrusion. The octadecyl rhodamine B chloride (R18), used as a lipophilic dye for collecting optical data, was made by Invitrogen purchased from Thermo Fisher Scientific-Waltham, MA. Dulbecco’s Modified Eagle Medium (DMEM) was used as a basal medium for cell growth and to produce pseudoparticles, along with Gibco Fetal Bovine Serum (FBS) and Gibco Penicillin-Streptomycin (10,000 U/mL) when indicated. TurboFect, Lipofectamine 2000, and Gibco Opti-MEM were purchased through Life Technologies Thermo Fisher and were used for the necessary transfection protocols described later in this section. Corning Trypsin 1×, 0.25% Trypsin purchased through VWR, 0.53 mM EDTA in HBSS [-] calcium, magnesium was used as the enzymatic agent during passaging. 4-(2-Hydroxyethyl)piperazine-1-ethanesulfonic acid (HEPES), dithiothreitol (DTT), and formaldehyde solution, used for the preparation of the blebs, were all purchased from MilliporeSigma. VWR 25 mm × 25 mm glass coverslips were used for the preparation of the supported lipid bilayers and as solid supports for the collection of optical data. The Piranha wash consisted of sulfuric acid (95-98%, VWR) and hydrogen peroxide (50 wt. % solution, Krackler Scientific). PEDOT:PSS (PH 1000) was purchased from Ossila (Sheffield, UK), (3-Glycidyloxypropyl)trimethoxysilane (GOPS) was purchased from MilliporeSigma.

### Buffers and other solutions

GPMV Buffer A: 2 mM CaCl_2_, 10 mM HEPES, 150 mM NaCl, pH 7.4 GPMV Buffer B: 2 mM CaCl_2_, 10 mM HEPES, 150 mM NaCl, 25 mM formaldehyde, 2 mM DTT pH 7.4 Reaction Buffer A: 137 mM NaCl, 2.7 mM KCl, 10 mM Na_2_HPO_4_, 1.8 mM KH_2_HPO_4_, pH 7.4 Reaction Buffer B: 137 mM NaCl, 2.7 mM KCl, 10 mM Na_2_HPO_4_, 1.8 mM KH_2_HPO_4_, pH 5.8 C-DMEM: DMEM, 10 % (v/v) FBS, Penicillin-Streptomycin (200 units/mL and 200 ug/mL) F-DMEM: DMEM, 10 % (v/v) FBS

### Cell Culture

African green monkey kidney cells (Vero E6) from ATCC, TMPRSS2 enhanced Vero E6 from the JCRB Cell Bank, and Human embryonic kidney cells (HEK-293T) from ATCC were maintained in C-DMEM at 37 °C in an incubator containing 5% CO_2_ and 95% air. All cells were passaged upon reaching 80-95% confluency by first washing the cells with Dulbecco’s phosphate-buffered saline (DPBS) and then enzymatically releasing them from the flasks using Trypsin EDTA 1x. Confluency was monitored using bright-field microscopy.

### GPMV (‘bleb’) preparation

GMPVs were prepared using previously established methodologies aimed to produce free GMPVs from attached cells. Once the cells have achieved >90% confluency, in preparation for blebbing, the cells were washed with GMPV buffer A (3x). Freshly prepared GPMV Buffer B was then added to the plate and incubated at 37 °C for 2 hours. Both GPMV Buffer A and GPMV Buffer B contain small amounts of CaCl_2_, as calcium has been found to be crucial for promoting an optimal fusion environment^47,48^. The buffer, now containing the GPMVs, was decanted into a conical tube and incubated on ice for 45 minutes. Post incubation the top 80% of the solution was collected, and the bottom 20% was disposed. The GMPVs were characterized using dynamic light scattering using a Malvern Panalytical Enigma Business Park, Grovewood Road Malvern, WR14 1XZ, UK Zetasizer MAL1026438 and NanoCyte. New GMPVs were prepared every two weeks to ensure that maximum protein activity was maintained.

### Preparation of Pseudotyped Particles

Human embryonic kidney cells HEK293 cells were seeded on 6-well plates with 2 mLs of C-DMEM solution per well. The cell density typically reached ∼50% confluence prior to proceeding to the next step. Transfection was performed with three plasmids encoding for the different proteins required to form pseudotyped particles: the envelope glycoprotein, MLV gag and pol proteins, and luciferase reporter. The total amount of DNA per well was 1 ug with 300 ng of gagpol, 400 ng of luciferase reporter, and 300 ng of the envelope protein (all sequences encoding for the genes can be found in Supplementary Table 1). First the plasmids encoding for gagpol and luciferace were combined and incubated at room temperature for five minutes. For a 50 mL solution, 1.25 mLs of optimem and 1.4 mLs of polyethyleneimine (PEI) were added to a 50 mL falcon tube. The envelope proteins were added to the tube as well, appropriately scaling the amount to the 50 mL total volume. The envelope proteins were either SARS-CoV2 spike protein, vesicular stomatitis virus (VSV) G glycoprotein, or a negative control that lacked any enveloped glycoproteins (Δenv). The backbone proteins (gagpol) were then added to the same tube and incubated at room temperature for 20 minutes. F-DMEM was added to a final volume of 50 mLs after the incubation. The C-DMEM was aspirated from the HEK293 cells and washed with F-DMEM prior to adding the transfection mixture. The F-DMEM mixture was then added, where each well on the plate contained a final volume of 2 mLs, and incubated for 48 hours at 37°C. By the end of the incubation period, the cells typically changed color to orange, being careful not to over-incubate (resulting in yellow color). The supernatant was collected from the wells and placed into 50 mL falcon tubes. These tubes were centrifuged for 7 minutes at 290 xg at 4°C. Being careful not to disturb the bottom of the tubes, the supernatant was, once again, recovered and filtered through a 0.22 μm syringe filter. To ensure longevity of the samples, 1 mL aliquots were frozen and stored at –80 °C until needed for use.

### Pseudotyped Particle transduction (infectivity) assay

Spike-containing viral pseudoparticles (VPPs_spike_) were produced as mimics of SARS-CoV-2 infectious virions using previously established methodologies^22^. The backbone of the VPPs consisted of a Murine Leukemia Virus (MLV)-gagpol and the viral envelope contained wtSARS-CoV2 Spike protein (WH1 strain), referred to as wt in the bar graph here. The interior cavity of the particles contained a luciferase reporter gene, which allowed for a straightforward method to test the transduction of the VPPs_spike_. In this assay, once the reporter gene was successfully delivered and integrated into the host cell’s genome, the transduced cells were quantified using a luciferase activity assay. To perform this assay, African green monkey kidney epithelial Vero-E6 cells were seeded in 24-well plates and incubated until 80-90% confluency was obtained. Each well was washed with 0.5 mLs of Dulbecco’s Phosphate Buffered Saline (DPBS) 3x, inoculated with 0.2 mLs of undiluted pseudovirus particle solution, and incubated at 37 °C for 1.5 hours while agitating on a rocker. After the first incubation period was complete, 0.2 mLs of C-DMEM were added and incubated at 37 °C for 72 hours. The infectivity was assessed using previously reported luciferase assay. Briefly, the luciferase substrate and 5× Promega lysis buffer were thawed. The buffer was diluted with sterile water and added to the cells for lysis. For a most effective lysis, the cells went through several freeze thaw cycles, being transferred from –80 °C to room temperature 3×. After the last thaw cycle, 10 μL of lysate and 20 μL of Luciferin were combined in an eppendorf tube and analyzed using a Promega (Durham, NC) GlowMax 20/20 luminometer.

### Transfection of plasmids containing SARS-CoV2 Spike

Typically the SARS-CoV2 Spike was transfected into HEK293 cells. For a 10 cm petri dish 400 μL of Opti-MEM was combined with either 24 μL of Lipofectamine and incubated for five minutes at room temperature. In another tube, 8 μg of plasmid was added to 400 μL of Opti-MEM. The two tubes were combined and incubated further for 20 minutes at room temperature. Once the appropriate cells reached ∼70% confluency, they were washed with DPBS and the Opti-MEM solution, containing transfection reagent and the plasmid, was added directly to the cells. The cells were incubated at 37 °C for one hour, then 8 mLs of C-DMEM were added to the top of the cells as well. They continued to incubate at 37 °C for 12-16 hours before the next step.

### SLB formation on PEDOT:PSS surface

To form SLB with cell blebs on PEDOT:PSS surface, a simultaneous incubation of both blebs and fusogenic vesicles is applied to generate repeatable results. PEDOT:PSS coverslip/ electrode device were soaked in DI for over 24 hours prior to use. Cell blebs and POPC lipids were mixed and sonicated for 20 mins to induce fusion^49,50^ before adding onto a light oxygen plasma-treated (Harrick Plasma Inc., Ithaca NY, PDC-32G, 7.2 W, 350 Micron, 1 min) PEDOT:PSS surface. It is worth noting that the plasma condition needs to be tuned for each plasma cleaner, as weak treatment won’t provide sufficient hydrophilicity to rupture the blebs and vesicles, while too strong of a treatment will render the surface more negative and rough, making it challenging for the often negative native components to self assemble into a mobile SLB. The incubation time for SLB formation on PEDOT:PSS surface was 1 hour before excess materials were rinsed out with PBS buffer prior to further characterizations. The presence of native membrane components in SLB was verified using TIRF as shown in Fig. 2b.

### FRAP analysis

Prior to SLB formation described in the previous method, the blebs were sonicated for 30 minutes (kept under 25°C with ice pad) to incorporate the lipophilic dye octadecyl rhodamine B chloride (R18) into the blebs (1 μL of 0.5 mg/mL R18 into 100 μL of blebs). SLB formation proceeded as previously described. To verify formation of the SLB and confirm lipid mobility, an inverted Zeiss Axio Observer Z1 microscope was used with a 20× objective lens. A 20 μm diameter was bleached for 500 ms and the recovery was monitored for 30 minutes. The fluorescence intensity was recorded and normalized. The data was fit to a standard Bessel function and diffusion (*D*) was determined using the equation: *D* = *w*^2^ */ 4t*_1/2_, where *w* represents the width (diameter) of the bleach spot, and *t*_1/2_ is the time it took for the fluorescence to recover to half of the maximum intensity, and *D* is the determined diffusion measurement.

### TIRF microscopy

SLBs were prepared (without the R18 dye as previously described). The pseudoparticles were first labeled by sonicating with R18 dye (1 μL of 0.5 mg/mL for 100 μL of pseudo particle solution) for 30 minutes (kept under 25°C with ice pad). For this assay, the VPPs are labeled to a semi-quenched state (independently verified using a fluorimeter (Supplementary Fig. 10)), where the fluorescence intensity is adequate to observe the particles within the TIRF field of view (FOV) but not proportional to the extent of labeling. The excess dye was removed using a size-exclusion column or simply washed away when appropriate. TIRF measurements were performed on Zeiss Axio Observer.Z1 microscope using an α Plan-Apochromat 100x objective with a numerical aperture (NA) of 1.46. The samples were excited with a 561 nm laser and the angle of incidence was adjusted to ∼68° to insure an evanescent wave of 100 nm with total internal reflection. Prior to acquiring these images, we washed our experimental well with excessive buffer to remove any unbound particles and ensure that we were acquiring images of only those particles that were bound and not diffusing in/out of the FOV.

### Microelectrode fabrication

Gold contact pads were patterned on fused silica wafer using a standard photolithography procedure: exposure, develop, deposition, and lift-off^51^. A 200 nm of SiO_2_ insulating layer was then deposited ubiquitously on Au patterned wafer using plasma enhanced chemical vapor deposition (PECVD). A second layer of photolithography was applied to define the PEDOT:PSS electrode locations on the gold contact pad, followed by the reactive ion etching of SiO_2_ until it reached the gold surface. PEDOT:PSS mixed with 1 v/v % of GOPS was then spin-coated at 4000 rpm on both exposed gold contact and SiO_2_ insulating layer, followed by the annealing at 140°C for 30 mins to drive off all water. A third layer of photolithography was applied to remove the PEDOT:PSS spun on SiO_2_, taking advantage of the germanium (Ge) hard mask protocol previously reported^52^. The 100 nm thick protective Ge hard mask on PEDOT:PSS electrode was then removed by immersing in deionized water for 48 hours.

### EIS Measurement and Data Analysis

An Autolab PGSTAT302N potentiostat was used to conduct the EIS measurements. The frequency of applied sinusoidal voltage was swept from 10^6^ Hz to 1 Hz to capture the change in electrical signal at each step after the addition of biological materials. Prior to SLB formation, the PEDOT:PSS electrode baseline was measured and fitted into a RC circuit. Signals after SLB formation were fit to a RC(RC) circuit, where membrane resistance and capacitance were extracted. Vigorous rinsing with PBS buffer was done before each measurement at every step after SLB formation, 20 mins after adding VLPs or cell blebs and 30 mins after adding CatL.

## Data availability

The data supporting the findings of this study are available within the paper and supplementary information (Fig. 1-12 and Table 1).

## Acknowledgements

S.D. and R.O. acknowledge funding for this project, sponsored by the Defense Advanced Research Projects Agency (DARPA) Army Research Office and accomplished under Cooperative Agreement Number W911NF-18-2-0152. The views and conclusions contained in this document are those of the authors and should not be interpreted as representing the official policies, either expressed or implied, of DARPA or the Army Research Office or the U.S. Government. The U.S. Government is authorized to reproduce and distribute reprints for Government purposes notwithstanding any copyright notation herein. The fabrication of microelectrodes in this work was performed at the Cornell NanoScale Facility, a member of the National Nanotechnology Coordinated Infrastructure (NNCI), which is supported by the National Science Foundation (Grant NNCI-2025233). Zhongmou Chao and Susan Daniel acknowledge Smith Fellowship for Postdoctoral Innovation from Cornell University. We thank Juliana D. Carten for useful discussions and assistance with the editing of the final manuscript.

## Author information

Authors and Affiliations

Robert Frederick Smith School of Chemical and Biomolecular Engineering, Cornell University, 124 Olin Hall, Ithaca, NY 14853, USA

Zhongmou Chao, Ekaterina Selivanovitch, Ambika Pachaury & Susan Daniel

Department of Chemical Engineering and Biotechnology, University of Cambridge, Philippa Fawcett Dr., Cambridge CB3 0AS, UK

Konstantinos Kallitsis, Zixuan Lu & Róisín Owens

## Contributions

Conceptualization: Zhongmou Chao, Ekaterina Selivanovitch, Konstantinos Kallitsis, Zixuan Lu, Róisín Owens, Susan Daniel

Methodology: Zhongmou Chao, Ekaterina Selivanovitch

Investigation: Zhongmou Chao, Ekaterina Selivanovitch, Ambika Pachaury

Visualization: Zhongmou Chao, Ekaterina Selivanovitch

Supervision: Róisín Owens, Susan Daniel

Writing—original draft: Zhongmou Chao, Ekaterina Selivanovitch, Susan Daniel

Writing—review & editing: Zhongmou Chao, Ekaterina Selivanovitch, Konstantinos Kallitsis, Zixuan Lu, Ambika Pachaury, Róisín Owens, Susan Daniel

## Corresponding author

Correspondence to Susan Daniel.

## References

1 Domingo, E., García-Crespo, C., Lobo-Vega, R. & Perales, C. Mutation Rates, Mutation Frequencies, and Proofreading-Repair Activities in RNA Virus Genetics. Viruses 13, 1882 (2021).

2 Drake, J. W. & Holland, J. J. Mutation rates among RNA viruses. Proceedings of the National Academy of Sciences 96, 13910–13913, doi:doi:10.1073/pnas.96.24.13910 (1999).

3 Elena, S. F. & Sanjuan, R. Adaptive value of high mutation rates of RNA viruses: separating causes from consequences. J Virol 79, 11555–11558, doi:10.1128/JVI.79.18.11555-11558.2005 (2005).

4 Shahabipour, F. et al. Engineering organ-on-a-chip systems to model viral infections. Biofabrication 15, doi:10.1088/1758-5090/ac6538 (2023).

5 Si, L. et al. A human-airway-on-a-chip for the rapid identification of candidate antiviral therapeutics and prophylactics. Nat Biomed Eng 5, 815–829, doi:10.1038/s41551-021-00718-9 (2021).

6 Tan, J. et al. Biomimetic lung-on-a-chip to model virus infection and drug evaluation. Eur J Pharm Sci 180, 106329, doi:10.1016/j.ejps.2022.106329 (2023).

7 Jackson, C. B., Farzan, M., Chen, B. & Choe, H. Mechanisms of SARS-CoV-2 entry into cells. Nature Reviews Molecular Cell Biology 23, 3–20, doi:10.1038/s41580-021-00418-x (2022).

8 Nolan, T., Hands, R. E. & Bustin, S. A. Quantification of mRNA using real-time RT-PCR. Nature protocols 1, 1559–1582 (2006).

9 Amanat, F. et al. A serological assay to detect SARS-CoV-2 seroconversion in humans. Nature medicine 26, 1033–1036 (2020).

10 Diao, B. et al. Accuracy of a nucleocapsid protein antigen rapid test in the diagnosis of SARS-CoV-2 infection. Clinical Microbiology and Infection 27, 289. e281–289. e284 (2021).

11 Chen, Y. et al. Impact of SARS-CoV-2 variants on the analytical sensitivity of rRT-PCR assays. Journal of Clinical Microbiology 60, e02374–02321 (2022).

12 Zou, Y., Mason, M. G. & Botella, J. R. Evaluation and improvement of isothermal amplification methods for point-of-need plant disease diagnostics. PloS one 15, e0235216 (2020).

13 Gootenberg, J. S. et al. Nucleic acid detection with CRISPR-Cas13a/C2c2. Science 356, 438–442 (2017).

14 Yousefi, H. et al. Detection of SARS-CoV-2 viral particles using direct, reagent-free electrochemical sensing. Journal of the American Chemical Society 143, 1722–1727 (2021).

15 Ventura, B. D. et al. Colorimetric test for fast detection of SARS-CoV-2 in nasal and throat swabs. ACS sensors 5, 3043–3048 (2020).

16 Dziąbowska, K., Czaczyk, E. & Nidzworski, D. Detection methods of human and animal influenza virus—current trends. Biosensors 8, 94 (2018).

17 Zhou, F. et al. A Supported Lipid Bilayer-Based Lab-on-a-Chip Biosensor for the Rapid Electrical Screening of Coronavirus Drugs. ACS Sens 7, 2084–2092, doi:10.1021/acssensors.2c00970 (2022).

18 Wrapp, D. et al. Cryo-EM structure of the 2019-nCoV spike in the prefusion conformation. Science 367, 1260–1263, doi:doi:10.1126/science.abb2507 (2020).

19 Hoffmann, M. et al. SARS-CoV-2 Cell Entry Depends on ACE2 and TMPRSS2 and Is Blocked by a Clinically Proven Protease Inhibitor. Cell 181, 271–280.e278, 10.1016/j.cell.2020.02.052 (2020).

20 Shulla, A. et al. A Transmembrane Serine Protease Is Linked to the Severe Acute Respiratory Syndrome Coronavirus Receptor and Activates Virus Entry. Journal of Virology 85, 873–882, doi:doi:10.1128/jvi.02062-10 (2011).

21 Bayati, A., Kumar, R., Francis, V. & McPherson, P. S. SARS-CoV-2 infects cells after viral entry via clathrin-mediated endocytosis. Journal of Biological Chemistry 296, 100306, 10.1016/j.jbc.2021.100306 (2021).

22 Au-Millet, J. K., et al. Production of Pseudotyped Particles to Study Highly Pathogenic Coronaviruses in a Biosafety Level 2 Setting. JoVE, e59010, doi:doi:10.3791/59010 (2019).

23 Zhang, Y. et al. Supported Lipid Bilayer Assembly on PEDOT:PSS Films and Transistors. Advanced Functional Materials 26, 7304–7313, 10.1002/adfm.201602123 (2016).

24 Liu, H.-Y. et al. Self-Assembly of Mammalian-Cell Membranes on Bioelectronic Devices with Functional Transmembrane Proteins. Langmuir 36, 7325–7331, doi:10.1021/acs.langmuir.0c00804 (2020).

25 Zhao, Z., Spyropoulos, G. D., Cea, C., Gelinas, J. N. & Khodagholy, D. Ionic communication for implantable bioelectronics. Science Advances 8, eabm7851, doi:doi:10.1126/sciadv.abm7851 (2022).

26 Rochford, A. E. et al. Functional neurological restoration of amputated peripheral nerve using biohybrid regenerative bioelectronics. Science Advances 9, eadd8162, doi:doi:10.1126/sciadv.add8162 (2023).

27 Costello, D. A., Hsia, C.-Y., Millet, J. K., Porri, T. & Daniel, S. Membrane Fusion-Competent Virus-Like Proteoliposomes and Proteinaceous Supported Bilayers Made Directly from Cell Plasma Membranes. Langmuir 29, 6409–6419, doi:10.1021/la400861u (2013).

28 Costello, D. A., Millet, J. K., Hsia, C.-Y., Whittaker, G. R. & Daniel, S. Single particle assay of coronavirus membrane fusion with proteinaceous receptor-embedded supported bilayers. Biomaterials 34, 7895–7904, 10.1016/j.biomaterials.2013.06.034 (2013).

29 Ogando, N. S. et al. SARS-coronavirus-2 replication in Vero E6 cells: replication kinetics, rapid adaptation and cytopathology. Journal of General Virology 101, 925–940, 10.1099/jgv.0.001453 (2020).

30 Matsuyama, S. et al. Enhanced isolation of SARS-CoV-2 by TMPRSS2-expressing cells. Proceedings of the National Academy of Sciences 117, 7001–7003, doi:doi:10.1073/pnas.2002589117 (2020).

31 Proctor, C. M., Rivnay, J. & Malliaras, G. G. Understanding volumetric capacitance in conducting polymers. Journal of Polymer Science Part B: Polymer Physics 54, 1433–1436, 10.1002/polb.24038 (2016).

32 Manzer, Z. A. et al. Cell-Free Synthesis Goes Electric: Dual Optical and Electronic Biosensor via Direct Channel Integration into a Supported Membrane Electrode. ACS Synthetic Biology 12, 502–510, doi:10.1021/acssynbio.2c00531 (2023).

33 Tang, T. et al. Functional Infectious Nanoparticle Detector: Finding Viruses by Detecting Their Host Entry Functions Using Organic Bioelectronic Devices. ACS Nano 15, 18142–18152, doi:10.1021/acsnano.1c06813 (2021).

34 Lubrano, C., Matrone, G. M., Iaconis, G. & Santoro, F. New Frontiers for Selective Biosensing with Biomembrane-Based Organic Transistors. ACS Nano 14, 12271–12280, doi:10.1021/acsnano.0c07053 (2020).

35 Lindholm-Sethson, B. Electrochemistry at Ultrathin Organic Films at Planar Gold Electrodes. Langmuir 12, 3305–3314, doi:10.1021/la951026k (1996).

36 Gritsch, S., Nollert, P., Jähnig, F. & Sackmann, E. Impedance Spectroscopy of Porin and Gramicidin Pores Reconstituted into Supported Lipid Bilayers on Indium−Tin-Oxide Electrodes. Langmuir 14, 3118–3125, doi:10.1021/la9710381 (1998).

37 Lowther, J. et al. The Importance of pH in Regulating the Function of the Fasciola hepatica Cathepsin L1 Cysteine Protease. PLOS Neglected Tropical Diseases 3, e369, doi:10.1371/journal.pntd.0000369 (2009).

38 Simmons, G. et al. Inhibitors of cathepsin L prevent severe acute respiratory syndrome coronavirus entry. Proceedings of the National Academy of Sciences 102, 11876–11881, doi:doi:10.1073/pnas.0505577102 (2005).

39 Park, S. B. et al. SARS-CoV-2 omicron variants harbor spike protein mutations responsible for their attenuated fusogenic phenotype. Communications Biology 6, 556, doi:10.1038/s42003-023-04923-x (2023).

40 Suzuki, R. et al. Attenuated fusogenicity and pathogenicity of SARS-CoV-2 Omicron variant. Nature 603, 700–705, doi:10.1038/s41586-022-04462-1 (2022).

41 Wang, X.-J. et al. Neutralization sensitivity, fusogenicity, and infectivity of Omicron subvariants. Genome Medicine 14, 146, doi:10.1186/s13073-022-01151-6 (2022).

42 Hu, B. et al. Spike mutations contributing to the altered entry preference of SARS-CoV-2 omicron BA.1 and BA.2. Emerging Microbes & Infections 11, 2275–2287, doi:10.1080/22221751.2022.2117098 (2022).

43 Meng, B. et al. Altered TMPRSS2 usage by SARS-CoV-2 Omicron impacts infectivity and fusogenicity. Nature 603, 706–714, doi:10.1038/s41586-022-04474-x (2022).

44 Willett, B. J. et al. SARS-CoV-2 Omicron is an immune escape variant with an altered cell entry pathway. Nature Microbiology 7, 1161–1179, doi:10.1038/s41564-022-01143-7 (2022).

45. Mizuki, Y. et al. SARS-CoV-2 Omicron spike H655Y mutation is responsible for enhancement of the endosomal entry pathway and reduction of cell surface entry pathways. bioRxiv, 2022.2003.2021.485084, doi:10.1101/2022.03.21.485084 (2022).

46 Kimura, I. et al. Virological characteristics of the SARS-CoV-2 Omicron BA.2 subvariants, including BA.4 and BA.5. Cell 185, 3992–4007.e3916, 10.1016/j.cell.2022.09.018 (2022).

47 Straus, M. R. et al. Ca^2+^ Ions Promote Fusion of Middle East Respiratory Syndrome Coronavirus with Host Cells and Increase Infectivity. Journal of Virology 94, 10.1128/jvi.00426-00420, doi:10.1128/jvi.00426-20 (2020).

48 Lai, A. L., Millet, J. K., Daniel, S., Freed, J. H. & Whittaker, G. R. The SARS-CoV Fusion Peptide Forms an Extended Bipartite Fusion Platform that Perturbs Membrane Order in a Calcium-Dependent Manner. Journal of Molecular Biology 429, 3875–3892, 10.1016/j.jmb.2017.10.017 (2017).

49 Pace, H. et al. Preserved Transmembrane Protein Mobility in Polymer-Supported Lipid Bilayers Derived from Cell Membranes. Analytical Chemistry 87, 9194–9203, doi:10.1021/acs.analchem.5b01449 (2015).

50 Thorsteinsson, K., Olsén, E., Schmidt, E., Pace, H. & Bally, M. FRET-Based Assay for the Quantification of Extracellular Vesicles and Other Vesicles of Complex Composition. Analytical Chemistry 92, 15336–15343, doi:10.1021/acs.analchem.0c02271 (2020).

51 Thompson, L. F. in Introduction to Microlithography Vol. 219 ACS Symposium Series Ch. 1, 1-13 (AMERICAN CHEMICAL SOCIETY, 1983).

52 Thiburce, Q., Melosh, N. & Salleo, A. Wafer-scale microfabrication of flexible organic electrochemical transistors. Flexible and Printed Electronics 7, 034001, doi:10.1088/2058-8585/ac808a (2022).

