## Supplemental Information for "Recreating the Biological Steps of Viral Infection on a Bioelectronic Platform to Profile Viral Variants of Concern"

Zhongmou Chao *et al.*

**This PDF file includes:**

Supplementary Text
Figs. S1 to S12
Table S1

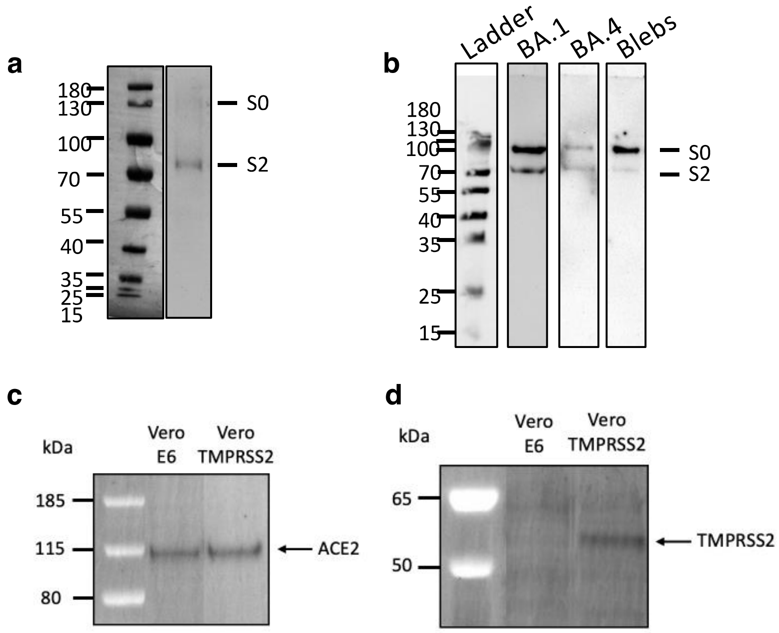


**Figure S1. Validation of Spike incorporation into SARS-CoV-2 pseudoparticles and ACE2 and TMPRSS2 in cells using Western Blots**. a) Representative western blot of pseudoparticles (VPPs) containing Spike (Wuhan-Hu-1). Pseudoparticles were resolved using gel electrophoresis, transferred, and then immuno-stained to visualize the SARS- CoV-2 S2 domain using a Spike rabbit polyclonal antibody (Sino Biological). b) Representative western blot of the ladder (lane 1) and pseudoparticles containing Omicron BA.1 (lane 2) and Omicron BA.4, (lane 3). Lane 4 contains blebs derived from HEK 293T cells transfected with wtSpike (Wuhan-Hu-1), which were used in our reverse configuration experiments where the SLB modeled the virus surface, while the blebs represented the cell surface. All the variant VPPs in (b) contained both cleaved (S2) and uncleaved (S0) Spike constructs, which potentially contributed to the loss in fusogenicity we observed on our platform compared to the wtSpike VPPs (a). A chemiluminescent western blot detection method was applied, using HRP-conjugated antibody coupled with enhanced chemiluminescent substrates. As this method is semi-quantitative, these western blots show only the presence or absence of a protein. To analyze the cell surface expression of receptor c) ACE2 and d) protease TMPRSS2 in VeroE6 (ATCC # CRL-1586) and Vero/TMPRSS2 (JCRB # 1818) cell lines, cells were were treated with 400 ul of biotin buffer (250 ug/ml ThermoFisher Sulfo-NHS-SS-Biotin in PBS) to label the surface proteins with biotin. Afterwards 50 mM glycine in PBS was then added for 30 min to the cells. Subsequently, the cells were lysed using a lysis buffer (0.1% TritonX in 1X TBS, 1 Complete Protease Inhibitor tablet from Sigma) and centrifuged at 14,000 rpm for 10 min at 4C. The supernatant was then added to 40 ul of Streptavidin beads (Pierce Thermofisher). Overnight incubation of the supernatant with the streptavidin beads at 4C ensured that the biotinylated proteins were bound to the beads. c) ACE2 was detected using human anti-rabbit polyclonal primary antibody (Cell Signaling Technology Cat # 4355S) and AlexaFluor 488 goat anti-rabbit secondary antibody. d) TMPRSS2 was detected using human anti-rabbit polyclonal primary antibody (NovusBio Cat # NBP2-38263) in blocking buffer and AlexaFluor 488 goat anti-rabbit secondary antibody in blocking buffer.

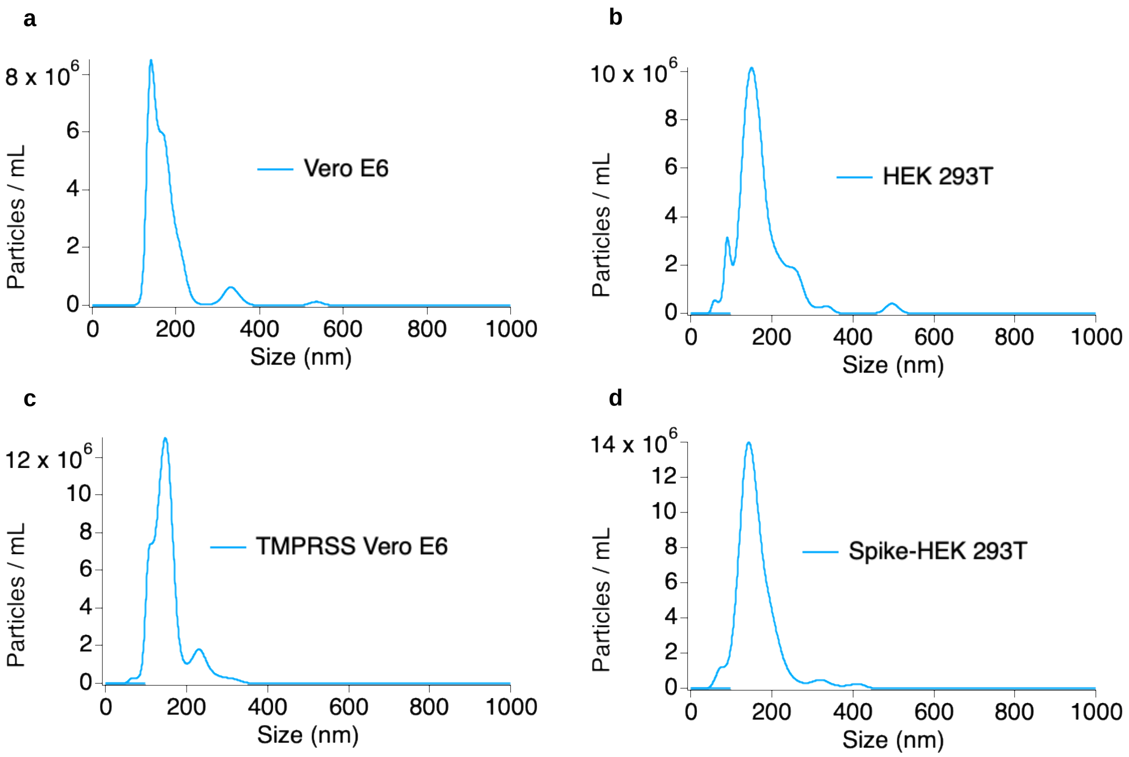


**Fig. S2. Bleb characterization using NanoSight Nanoparticle Tracking Analyzer**. As the blebs degrade, the components aggregate and form precipitates. Therefore, we used the measured particle size and count to assess bleb integrity. We measured the following samples: a) Vero E6, b) HEK 293T, c) TMPRSS2 Vero E6, and d) wtSpike-transfected HEK 293T. These are representative measurements taken for each of the bleb types.

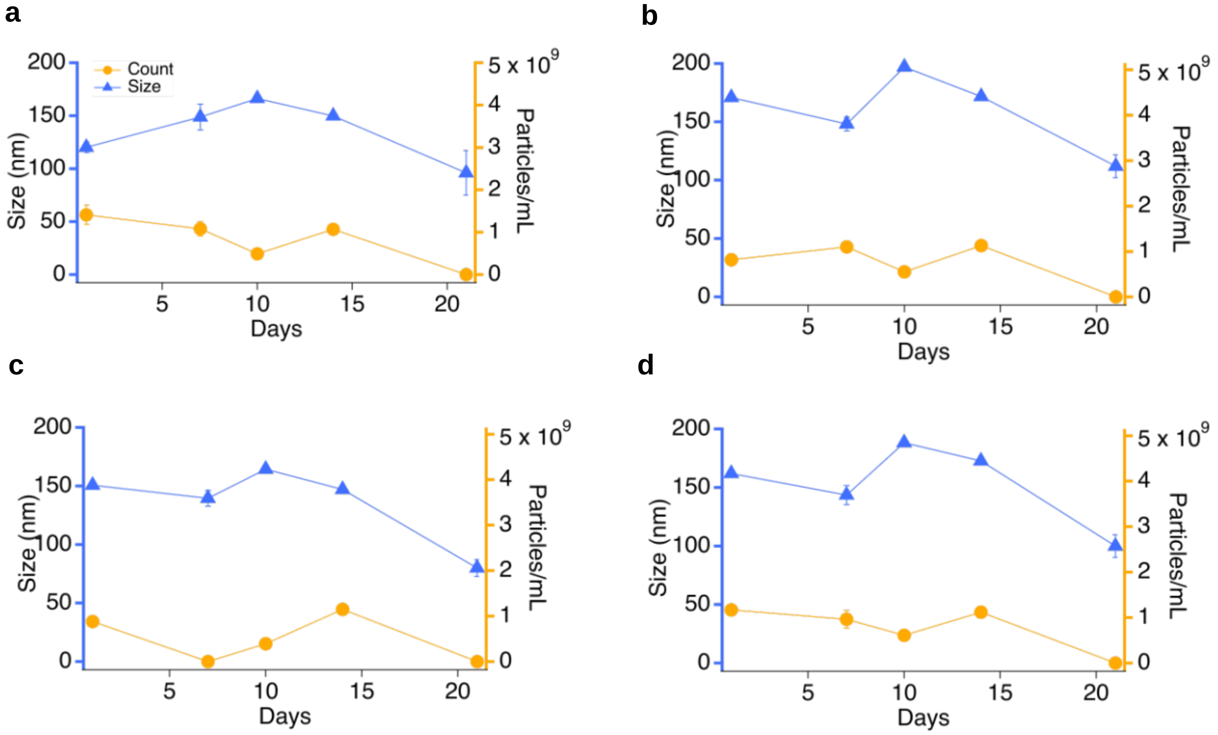


**Fig. S3. Data analysis of values obtained from Dynamic Light Scattering over 21 days**. The hydrodynamic radii were determined for blebs derived for the four cells lines used for this work, which include (a) Vero E6 cells, (b) HEK293-T cells, (c) Vero E6 TMPRSS2, and (d) wtSpike-transfected HEK293T cells. The sizes are depicted in the blue traces over the course of a three-week period. The average sizes remained consistent over the initial two-week period but decreased on the third week. These data aligned with the particle counts determined for each of the four bleb types (orange traces) over the same period. This was suggestive of particle degradation and was used as a guideline for experiments completed on the same set of blebs, which were confined to a two-week period. Error bars were calculated from the standard deviation (n = 3).

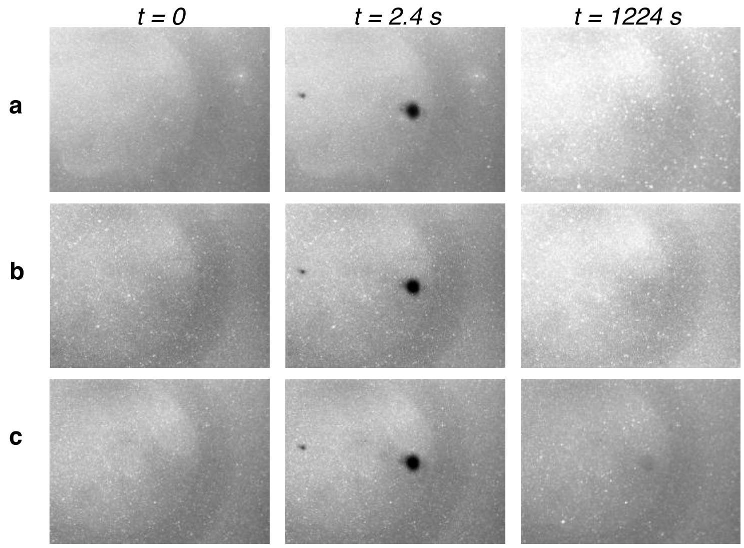


**Fig S4. Fluorescence Recovery After Photobleaching (FRAP) data collected using supported lipid bilayers (SLBs) assembled using Vero E6 cell-derived blebs, done in triplicate with recovery curves for each respective sample**. Qualitative analysis of these data indicates that the bleach spot recovers suggesting we indeed have a mobile bilayer. Quantitative analysis of these data can be found in **Fig S6**.

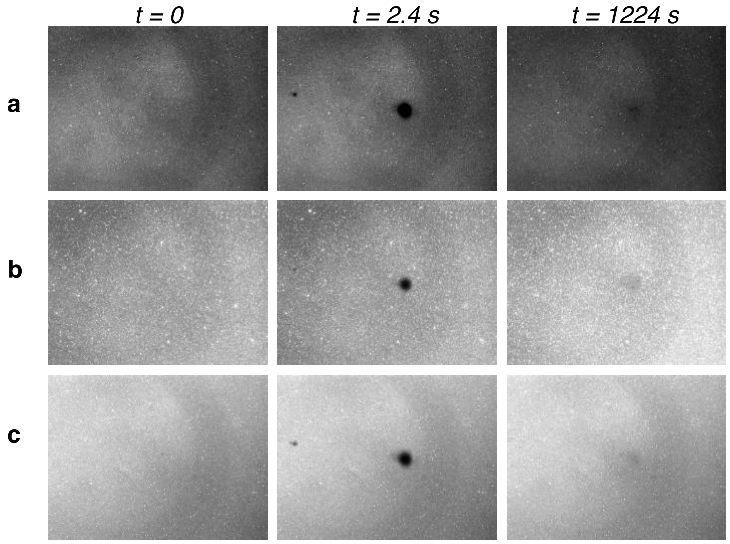


**Fig S5. Fluorescence Recovery After Photobleaching (FRAP) data collected using supported lipid bilayers (SLBs) assembled using Vero E6 TMPRSS2, done in triplicate with recovery curves for each respective sample**. Like the Vero E6 data presented in Fig S5, qualitative analysis of these data indicates a mobile bilayer. Quantitative analysis of these data can be found in Fig S6.

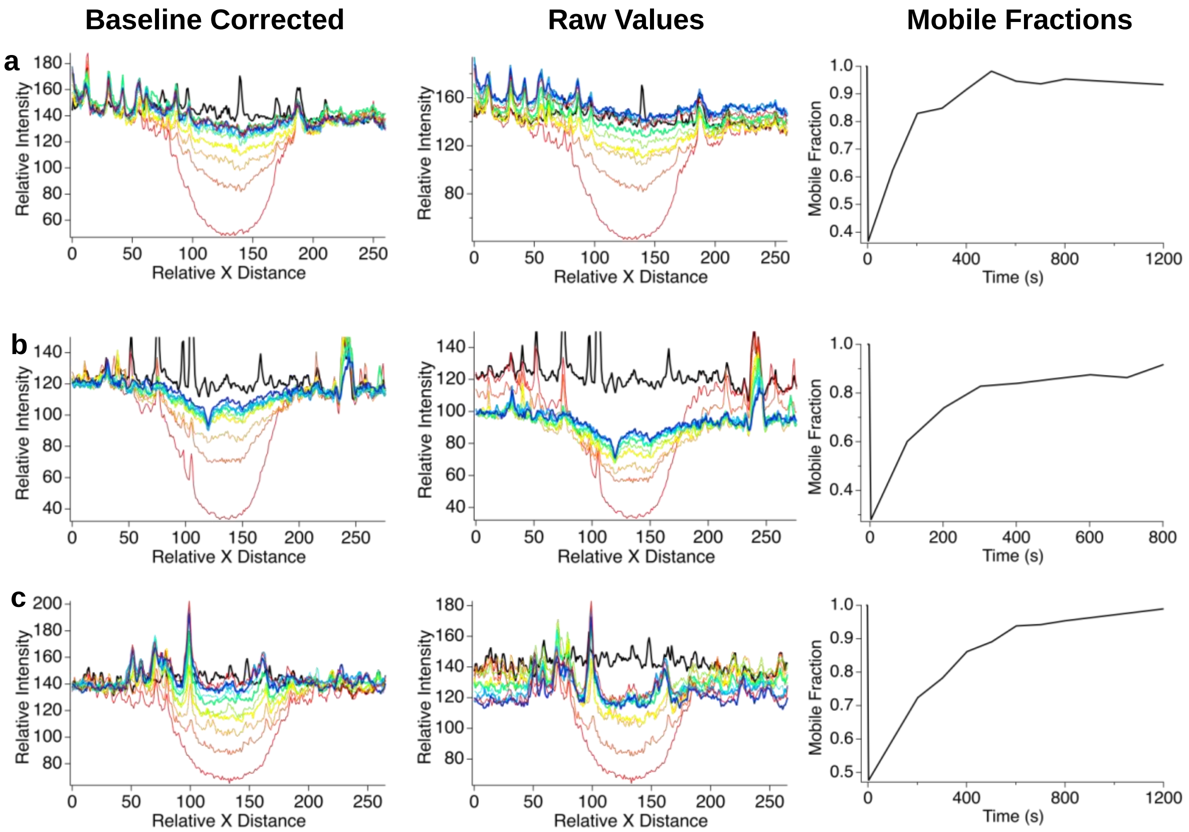


**Fig S6. Intensity line profiles of images acquired during FRAP experiments**. To account for baseline shifts during FRAP data acquisitions, we performed an intensity line plot analysis of the bleach spots before bleaching (bold black trace) and at several time points after including after ~ 1,200 s (bold blue trace- the last time point). The graphs on the left represent data after baseline corrections were (Baseline Corrected), and the ones on the right represent raw values acquired from the image analysis (Raw Values). The relative X distances (x-axis) represent the line scan across the image, with ~ 0 - 75 and ~ 175 - 250 representing the background (baseline) and 75 - 175 representing the bleach spot. The relative intensity is found on the y-axis and as can be seen on the graphs, the intensity dramatically drops and begins recovering immediately. This suggests that we indeed have a mobile bilayer. We used this data to calculate the true mobile fraction, rather than the apparent mobile fractions calculated prior to baseline correction (Fig S5, S6, S9). (a) Vero E6 Bleb-SLBs have a mobile fraction of 0.93 and diffusion (D) of 0.18 $\mu m^2 s^{-1}$, (b) TMPRSS2Vero E6 Bleb-SLBs have a mobile fraction of 0.92 and D of 0.16 $\mu m^2 s^{-1}$ (c) Spike-transfected HEK293T Bleb-SLBs have a mobile fraction of 0.99 and D of 0.20 $\mu m^2 s^{-1}$.

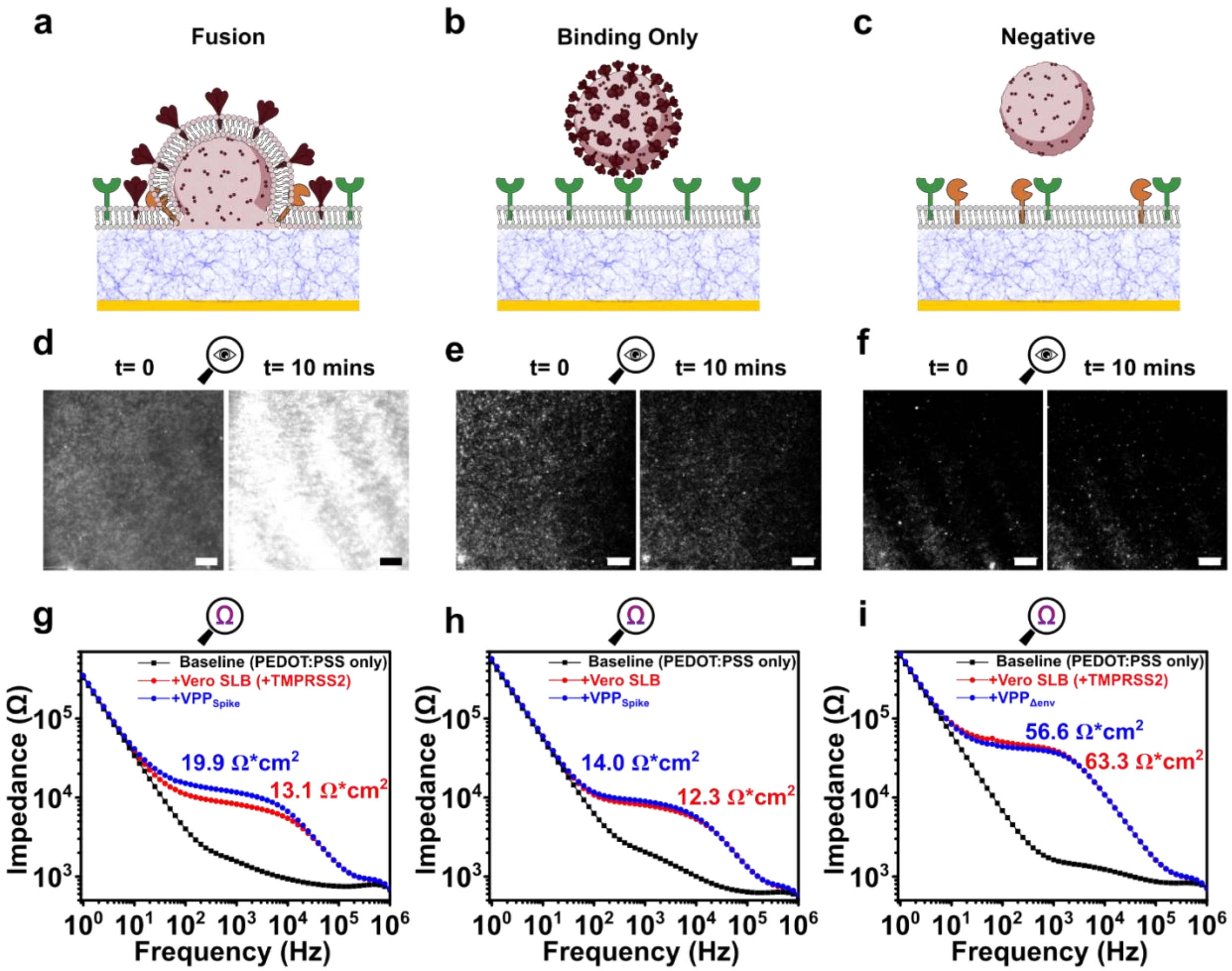


**Fig. S7. Early entry pathway recapitulated on host cell derived SLB**. (**a**, **d**, **g**) The experimental group consisted of $VPP_{Spike}$ and SLB containing ACE2 (green) and TMPRSS2 (yellow) where signals are characteristic of fusion events, with (d) showing the TIRF field of view (FOV) and changes in fluorescence after 10 minutes and (g) showing the changes in SLB membrane resistance in the equivalent electrical circuit scenario; (**b**, **e**, **h**) show one of the control groups where only signals coinciding to binding events are generated. This group consisted of $VPP_{Spike}$ and SLBs containing only ACE2, (e) shows the TIRF data and (h) impedance data; (**c**, **f**, **i**) show a negative control group where neither binding nor fusion are observed since $VPP_{\Delta env}$ were used, (f) shows the TIRF data and (i) shows impedance data; All scale bars represent 10 μm.

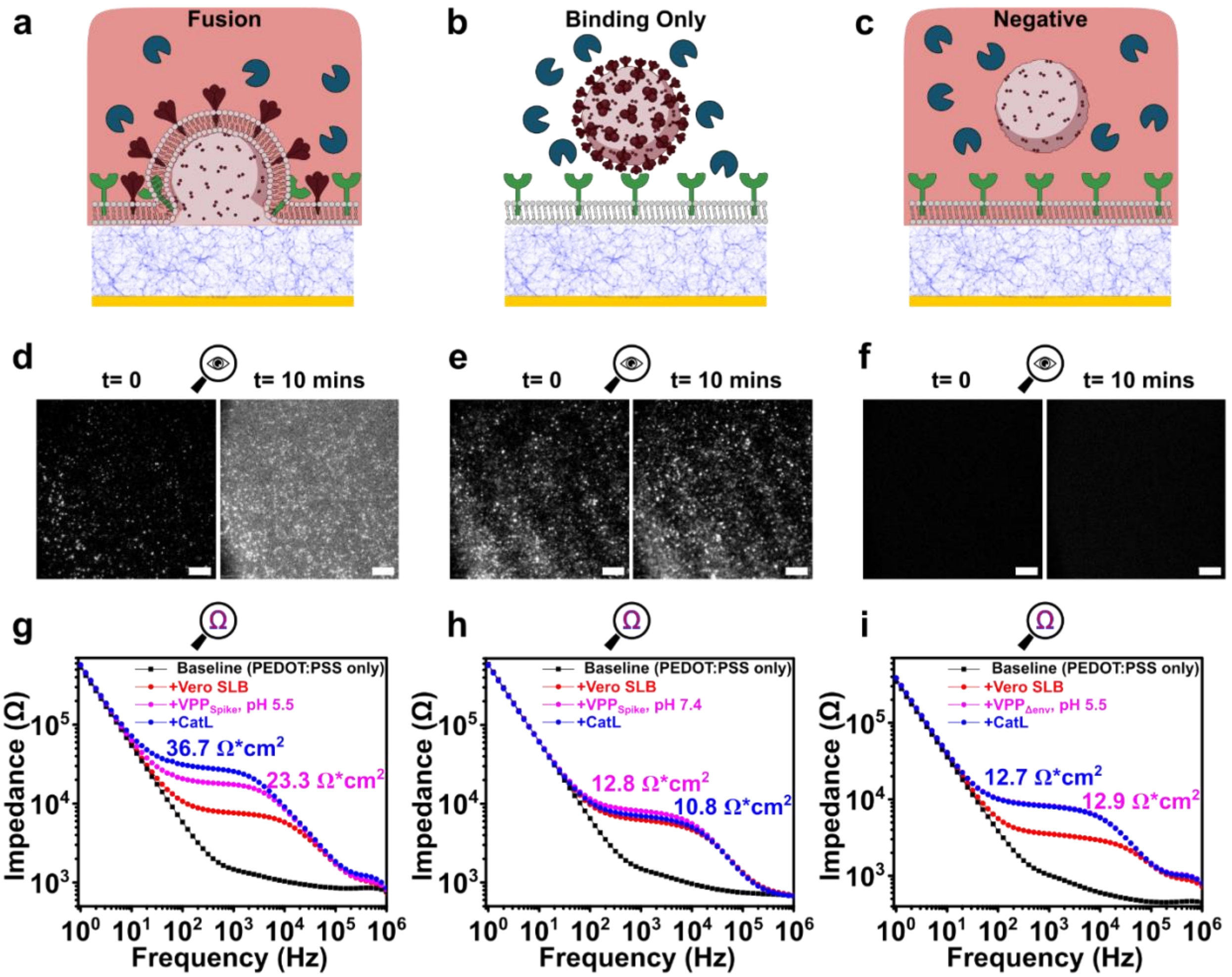


**Fig. S8. Late entry pathway recapitulated on host cell derived SLB**. (**a**, **d**, **g**) the experimental group consisted of VPP$_{Spike}$ and SLB containing ACE2 (green) and CatL (navy), where signals are characteristic of fusion events (note: pink droplet = acidic environment), (d) shows the TIRF field of view (FOV) and changes in fluorescence after 10 minutes while (g) shows the SLB resistance change upon fusion after the addition of CatL at pH = 5.5; (**b**, **e**, **h**) show one of the control groups where only signals coinciding to binding events are generated, this group consisted of VPP$_{Spike}$ and SLBs containing ACE2 but CatL was added into a nonacidic buffer environment (pH = 7.4), (e) shows the TIRF data and (h) the impedance data; (**c**, **f**, **i**) show a negative control group where neither binding nor fusion are observed since VPP$_{\Delta env}$ were used, (f) shows the TIRF data and (i) shows SLB resistance change upon the addition of CatL at pH = 5.5; All scale bars represent 10 μm.

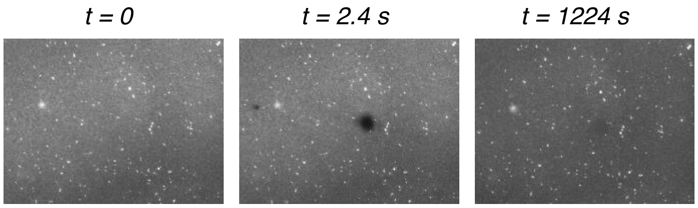


**Fig S9. Fluorescence Recovery After Photobleaching (FRAP) data collected using supported lipid bilayers (SLBs) assembled using wtSpike-transfected HEK 293T cells**. Qualitative analysis of these data indicates that the bleach spot recovers suggesting we indeed have a mobile bilayer. Quantitative analysis of these data can be found in Fig S7.

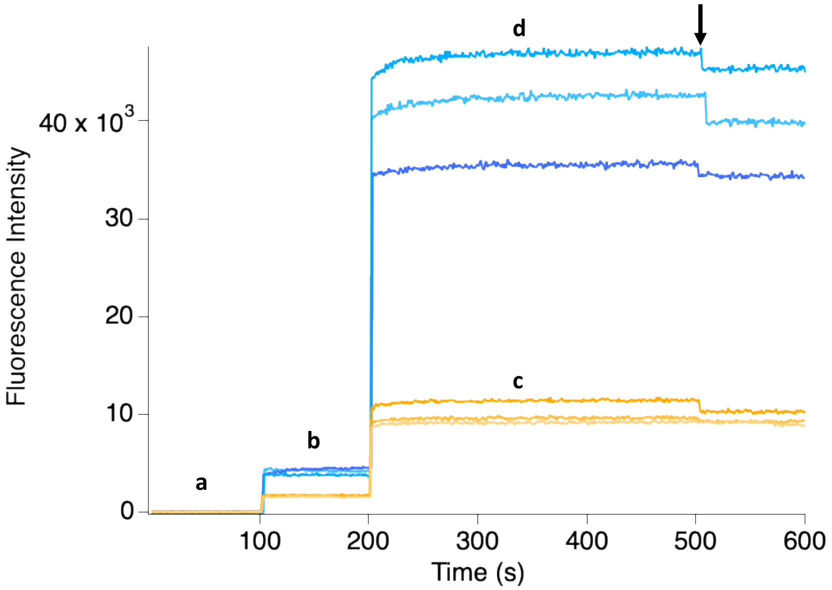


**Fig S9. Using fluorescence signal to optimize R18 labeling of SARS-CoV-S pseudoparticles to achieve semi-quenched state**. ~ $10^9$ Wuhan-Hu-1 Spike pseudoparticles/mL were labeled with two different R18 concentrations: 500 ng (orange) versus 5,000 ng (blue) and the samples were placed in a fluorimeter to compare their fluorescent traces. a) represents the first 100 seconds of the trace and is therefore the background signal; b) indicates when the pseudoparticles were added at t =100 seconds and shows the corresponding increase in fluorescence. At t = 200 seconds, 50 µLs of a 10 % (v/v) Triton-X 100 solution were added to the 1 mL pseudoparticle samples. The results indicated that labeling with 500ng of R18 did not result in pseudoparticles being in a quenched state, as evidenced by the small increase in signal upon the addition of detergent (c). Conversely, the fluorophores were in a semi-quenched state when 10-fold more R18 was used to label the pseudoparticles (d). This is indicated by the dramatic increase in signal upon Triton-X 100 addition, in comparison to the low but detectable signal prior to detergent addition. The dip in fluorescence observed at t = 500 seconds was due to additional detergent being added to confirm dequenching was complete (black arrow).

**Table S1. Sequences for the three Spike variants used for this study including Wuhan-Hu-1 Spike, Omicron BA.1, and Omicron BA.4, Gag-Pol, and Luciferase**.

| Name | Sequence |
|---|---|
| **Spike protein** | atgttcctgctgaccacaaagcggacaatgttcgtgtttctggtgctgctgcctctggtgagctcccagtgcgtgaacctg<br>accacaagaacccagctgccccctgcctataccaattccttcacacggggcgtgtactatcccgacaaggtgtttagatc<br>tagcgtgctgcactccacacaggatctgtttctgcctttcttttctaacgtgacctggttccacgccatccacgtgagcggc<br>accaatggcacaaagaggttcgacaatccagtgctgccctttaacgatggcgtgtacttcgcctccaccgagaagtcta<br>acatcatccgcggctggatctttggcaccacactggacagcaagacacagtccctgctgatcgtgaacaatgccacca<br>acgtggtgatcaaggtgtgcgagttccagttttgtaatgatccattcctgggcgtgtactatcacaagaacaataagtcttg<br>gatggagagcgagtttcgggtgtattcctctgccaacaattgcacatttgagtacgtgtcccagcccttcctgatggacct<br>ggagggcaagcagggcaatttcaagaacctgcgggagttcgtgtttaagaatatcgatggctacttcaagatctactcca<br>agcacaccccaatcaacctggtgagagacctgccacagggattctctgccctggagccactggtggatctgcccatcg<br>gcatcaacatcacccggtttcagacactgctggccctgcacagaagctacctgacaccaggcgacagctcctctggat<br>ggaccgcaggagctgccgcctactatgtgggctatctgcagccccggaccttcctgctgaagtacaacgagaatggca<br>ccatcacagacgcagtggattgcgccctggacccccctgtctgagaccaagtgtacactgaagagctttaccgtggaga<br>agggcatctatcagacaagcaatttcagggtgcagcctaccgagtccatcgtgcgctttcccaatatcacaaacctgtgc<br>ccttttggcgaggtgttcaacgccaccagattcgccagcgtgtacgcctggaataggaagcgcatctccaactgcgtgg<br>ccgactattctgtgctgtacaacagcgcctccttctctacctttaagtgctatggcgtgagccccacaaagctgaatgatct<br>gtgctttaccaacgtgtacgccgattccttcgtgatcaggggagacgaagtgaggcagatcgcaccaggacagacag<br>gaaagatcgcagactacaattataagctgcctgacgatttcaccggctgcgtgatcgcctggaactctaacaatctggat<br>agcaaagtgggcggcaactacaattatctgtaccggctgtttagaaagtctaatctgaagccattcgagcgggacatctc<br>cacagagatctaccaggccggctctacccccctgcaatggcgtggagggctttaactgttatttccctctgcagagctacg<br>gcttccagccaaccaacggcgtgggctatcagccctacagagtggtggtgctgtcttttgagctgctgcacgcacctgc<br>aacagtgtgcggcccaaagaagagcaccaatctggtgaagaacaagtgcgtgaacttcaacttcaacggactgaccg |

gcacaggcgtgctgaccgagtccaacaagaagttcctgccttttcagcagttcggcagggacatcgcagataccacag

acgccgtgcgcgaccctcagaccctggagatcctggacatcacaccatgctccttcggcggcgtgtctgtgatcacac

caggcaccaatacaagcaaccaggtggccgtgctgtatcaggacgtgaattgtaccgaggtgcccgtggcaatccac

gcagatcagctgacccctacatggcgggtgtactctaccggcagcaacgtgttccagacaagagccggatgcctgatc

ggagcagagcacgtgaacaatagctatgagtgcgacatccctatcggcgccggcatctgtgcctcctaccagaccca

gacaaactccccaaggagagccaggtctgtggccagccagtccatcatcgcctataccatgagcctgggcgccgaga

acagcgtggcctactccaacaattctatcgccatccctaccaacttcacaatctccgtgaccacagagatcctgccagtg

agcatgaccaagacatccgtggactgcacaatgtatatctgtggcgattccaccgagtgctctaacctgctgctgcagta

cggctctttttgtacccagctgaatcgcgccctgacaggaatcgcagtggagcaggacaagaacacacaggaggtgtt

cgcccaggtgaagcagatctacaagaccccacccatcaaggactttggcggcttcaacttcagccagatcctgcccga

tcctagcaagccatccaagaggtcttttatcgaggacctgctgttcaacaaggtgaccctggccgatgccggcttcatca

agcagtatggagattgcctgggagacatcgcagcccgcgacctgatctgtgcccagaagtttaatggcctgaccgtgct

gcctccactgctgacagatgagatgatcgcccagtacacatctgccctgctggccggcaccatcacaagcggatggac

cttcggcgcaggagccgccctgcagatcccctttgccatgcagatggcctatcggttcaacggcatcggcgtgaccca

gaatgtgctgtacgagaaccagaagctgatcgccaatcagtttaactccgccatcggcaagatccaggactctctgagc

tccacagccagcgccctgggaaagctgcaggatgtggtgaatcagaacgcccaggccctgaataccctggtgaagc

agctgtctagcaacttcggcgccatctcctctgtgctgaatgacatcctgagccggctggacaaggtggaggcagagg

tgcagatcgaccggctgatcacaggcagactgcagtccctgcagacctacgtgacacagcagctgatcagggcagca

gagatcagggcctctgccaatctggccgccaccaagatgagcgagtgcgtgctgggacagtccaagagggtggactt

ttgtggcaagggctatcacctgatgagcttcccacagtccgcccctcacggcgtggtgtttctgcacgtgacctacgtgc

cagcccaggagaagaacttcaccacagcaccagccatctgccacgatggaaaggcacactttcctagggagggcgt

gttcgtgagcaacggcacccactggtttgtgacacagcgcaatttctacgagccacagatcatcaccacagacaatacc

ttcgtgagcggcaactgtgatgtggtgatcggcatcgtgaacaataccgtgtatgatcctctgcagccagagctggactc

ttttaaggaggagctggataagtacttcaagaatcacaccagccccgacgtggatctgggcgacatctctggcatcaat

gccagcgtggtgaacatccagaaggagatcgacagactgaacgaggtggccaagaatctgaacgagtccctgatcg

atctgcaggagctgggcaagtatgagcagtacatcaagtggccctggtatatctggctgggcttcatcgccggcctgat
cgccatcgtgatggtgaccatcatgctgtgctgtatgacaagctgctgttcctgcctgaagggctgctgttcttgtggcag
ctgctgtaagtttgatgaggacgatagcgagcctgtgctgaagggcgtgaagctgcactacacctga

atgttcgtgtttctggtgctgctgcctctggtgtccagccagtgtgtgaacctgaccaccagaacacagctgcctccagc
ctacaccaacagctttaccagaggcgtgtactaccccgacaaggtgttcagatccagcgtgctgcactctacccaggac
ctgttcctgcctttcttcagcaacgtgacctggttccacgtgatcagcggcaccaatggcaccaagagattcgacaaccc
cgtgctgcccttcaacgacggggtgtactttgccagcatcgagaagtccaacatcatccgcggctggatcttcggcacc
acactggatagcaagacccagagcctgctgatcgtgaacaacgccaccaacgtggtcatcaaagtgtgcgagttccag
ttctgcaacgacccattcttcgaccacaagaacaacaagagctggatggaaagcgagttccgggtgtacagcagcgcc
aacaactgcaccttcgagtacgtgtcccagcctttcctgatggacctggaaggcaagcagggcaacttcaagaacctg
cgcgagttcgtgttcaagaacatcgacggctacttcaagatctacagcaagcacacccctatcatcgtgcgcgagcctg
aggatctgcctcagggcttttctgccctggaacctctggtggatctgcccatcggcatcaacatcacccggtttcagaca
ctgctggccctgcacagaagctacctgacacctggcgatagcagcagcggatggacagctggtgccgccgcttactat
**BA.1** gtgggctacctgcagcctagaaccttcctgctgaagtacaacgagaacggcaccatcaccgacgccgtggattgtgct
ctggatcccctgagcgagacaaagtgcaccctgaagtccttcaccgtggaaaagggcatctaccagaccagcaacttc
cgggtgcagcccaccgaatccatcgtgcggttccccaatatcaccaatctgtgcccctttgatgaggtgttcaatgccac
cagattcgccagcgtgtacgcctggaaccggaagagaatcagcaactgcgtggccgactactccgtgctgtacaatct
ggccccattctttaccttcaagtgctacggcgtgtcccctaccaagctgaacgacctgtgcttcaccaatgtgtacgccga
cagcttcgtgatccggggagatgaagtgcggcagattgcccctggacagaccggcaatatcgccgactacaactaca
agctgcccgacgacttcaccggctgtgtgatcgcctggaatagcaacaagctggacagcaaggtgtccggcaactac
aattacctgtaccggctgttccggaagtccaatctgaagcccttcgagcgggacatcagcaccgagatctatcaggccg
gcaacaagccctgtaatggcgtggccggcttcaactgctacttcccactgcggagctacagcttcagacccacatacg
gcgttggccaccagccttacagagtggtggtgctgtccttcgagctgctgcatgctcctgccacagtgtgcggccctaa
gaaaagcaccaacctcgtgaagaacaaatgcgtgaacttcaacttcaacggcctgaaaggcaccggcgtgctgaccg

agagcaacaagaagttcctgccattccagcagttcggccgggacattgccgataccacagatgctgtcagagatcccc

agacactggaaatcctggacatcacccccttgcagcttcggcggagtgtctgtgatcacccctggcaccaacaccagca

atcaggtggcagtgctgtaccagggcgtgaactgtacagaggtgccagtggccattcacgccgatcagctgacccccta

cttggcgggtgtactccacaggcagcaatgtgttccagaccagagccggctgtctgattggcgccgagtatgtgaaca

acagctacgagtgcgacatccccatcggagccggcatctgtgccagctaccagacacagaccaagagccacagacg

ggctagaagcgtggccagccagagcatcattgcctacacaatgtctctgggcgccgagaacagcgtggcctacagca

acaactctatcgctatccccaccaacttcaccatcagcgtgaccaccgagattctgcccgtgtccatgaccaagaccag

cgtggactgcaccatgtacatctgcggcgattccaccgagtgctccaacctgctgctgcagtacggcagcttctgcacc

cagctgaagagagagccctgacagggattgccgtggaacaggacaagaacacccaagaggtgttcgcccaagtgaagc

agatctacaagacccctcctatcaagtacttcggcgggttcaacttctcccagatcctgccagatcctagcaagcccagc

aagcggagcttcatcgaggacctgctgttcaacaaagtgacactggccgacgccggctttatcaagcagtatggcgatt

gcctgggcgacattgcagccagggatctgatttgcgcccagaagttcaagggcctgacagtgctgcctcctctgctgac

agatgagatgatcgcccagtacacaagcgccctgctggccggcacaatcacctctggatggacatttggagccggcg

ctgccctgcagatcccatttgctatgcagatggcctaccggttcaacggcatcggagtgacccagaatgtgctgtacga

gaaccagaagctgatcgccaaccagttcaacagcgccatcggcaagatccaggacagcctgagcagcacagcctct

gctctgggcaagctgcaggacgtggtcaaccataatgcccaggcactgaacaccctggtcaagcagctgtcctccaa

gttcggcgccatctctagcgtgctgaatgacatcttctccaggctggacaaggtggaagccgaggtgcagatcgacag

actgatcaccggaaggctgcagtccctgcagacctacgttacccagcagctgattagagccgccgagatcagagcca

gcgccaatctggctgccaccaagatgtctgagtgtgtgctgggccagagcaagagagtggacttttgcggcaagggct

accacctgatgagcttccctcagtctgctcctcacggcgtggtgtttctgcacgtgacatacgtgcccgctcaagagaag

aatttcaccaccgctccagccatctgccacgacggcaaagcccactttcctagagaaggcgtgttcgtgtccaacggca

cccattggttcgtgacccagcggaacttctacgagccccagatcatcaccaccgacaacaccttcgtgtctggcaactg

cgacgtcgtgatcggcattgtgaacaataccgtgtacgaccctctgcagcccgagctggactccttcaaagaggaactg

gataagtactttaagaaccacacaagccccgacgtggacctgggcgatatcagcggaatcaatgccagcgtcgtgaac

atccagaaagagatcgaccggctgaacgaggtggccaagaatctgaacgagagcctgatcgacctgcaagaactgg

ggaagtacgagcagtacatcaagtggccttggtacatctggctgggctttatcgccggactgattgccatcgtgatggtc
acaatcatgctgtgctgtatgaccagctgctgtagctgcctgaagggctgttgtagctgtggctcctgctgctga

**BA.4**

atgttcgtgtttctggttctcctgcccctggtgagcagccagtgtgttaatctgatcacccggacgcagagctatacaaata
gcttcaccagaggcgtgtactatcctgataaggtgttcagaagcagcgtgctgcacagcacacaagatctgttcctgcct
tttttcagcaatgtgacctggttccacgccatcagcggcaccaacggcaccaagcggtttgacaaccctgtgctgccttt
caacgatggggtgtacttcgcctctacagagaagagcaacatcatccggggctggatcttcggcaccaccctggattct
aagacccagagcttgctgatcgtgaacaatgctaccaacgtggtgatcaaagtgtgtgaattccagttctgcaacgaccc
ttttctggatgtgtactaccacaagaacaacaagtcttggatggaaagcgagttcagagtgtattcatctgccaacaactg
caccttcgagtacgtgtctcaacctttcctgatggacctggaaggcaagcagggcaacttcaagaaccttagagaattcg
tgttcaagaacatcgacggctacttcaagatctactctaagcacacacccatcaacctgggacgggacctcccccaagg
cttcagcgcccttgagcccctggtggacctgcctatcggcatcaacatcacccggttccagaccctgctggctctgcata
gaagctacctgaccccaggcgactctagcagcggctggaccgccggagccgccgcctactatgtgggctacctgca
acctagaactttcctgctcaagtacaatgagaatggcaccatcaccgacgccgtcgactgcgccctggatcctctgagc
gagacaaagtgcacactgaaaagtttcaccgtggaaaaaggcatctatcagacctctaactttagagtgcaacctaccg
agtcaatcgtgcggttccctaacatcaccaatctgtgtccttttgacgaggtgttcaacgctacaaggttcgccagcgtgt
acgcctggaaccggaaacggatctccaattgcgtggccgactacagcgtgctgtacaacttcgcccctttctttgccttc
aagtgctacggagtgtctccaacaaagctgaatgacctgtgcttcaccaatgtgtacgcagacagcttcgtgatcagag
gcaacgaggtgagccaaatcgcccccggccagacaggaaacattgccgattacaactacaagctacctgacgatttca
ccggctgcgtgatagcctggaactctaacaagctggatagcaaggtgggaggaaactacaactacagatacagactgt
tcagaaagtctaacctgaaaccttttgaaagagatatctctaccgagatctaccaggccggtaacaaaccgtgcaacgg
agtggccggcgtgaactgctactttccactgcagagctacggcttcagaccaacctacggcgttggccaccagccttac
cgggtggtggtgctgagcttcgagctgctgcacgcccctgccaccgtgtgcggacctaagaaatcgacaaacctggtg
aaaaacaagtgcgtgaattttaacttcaacggcctgacaggcacaggcgtgctgacagaaagtaacaaaaagttcctgc
ccttccagcagttcggaagagatatcgccgacaccacagatgccgtgcgggacccccagaccttggagatcctggac

atcacaccttgtagctttggcggcgtgagcgtcataaccccaggcacaaataccagcaaccaggtggccgtgctgtac

cagggcgtgaactgcaccgaggttcccgtggctattcacgccgaccagctgacacctacatggcgggtgtacagcac

cggctctaacgtgttccagaccagagccggctgcctgatcggagctgagtatgtgaacaacagctatgaatgtgacatc

cctatcggagctggcatttgtgccagctaccagacccagacgaaaagccaccgcagagccagaagcgtcgccagcc

agagtatcatcgcctacacgatgagcctgggcgcagagaactccgtggcctactccaacaactctatcgccatcccca

caaacttcactatctctgtcacaaccgaaattctgcccgtgagtatgaccaagaccagcgtcgactgcaccatgtacatct

gtggcgacagcacagagtgtagcaacctgctgctgcagtacgggagcttttgtacacagctgaagagagccctgacg

ggcatcgcagttgaacaggacaagaatacccaggaggtgttcgcccaggtgaagcagatttacaagacccctcctatt

aagtactttggcggattcaacttcagccagatcctgcctgaccctagcaagccttcgaagcggagcttcatcgaggacct

tctctttaacaaagtgacgctggccgacgccggcttcatcaagcagtacggcgactgcctgggcgacattgcagctag

agacctgatctgcgcccagaagtttaacggcctgaccgtgctgcctcctctgctgaccgacgaaatgatcgctcaatac

acaagcgccttactggccggcaccatcacttccggatggacattcggcgccggcgccgccctgcagattcctttcgcta

tgcagatggcttatcgcttcaacggcatcggcgtgacccagaacgtgctatacgagaaccagaagctgatcgccaatc

agtttaactcagctattggcaagatccaggattccctgtcctctacagccagcgccctgggtaaactgcaagatgtggtg

aaccacaacgctcaggccctgaacacactggtgaagcagctgagctccaagtttggcgccatcagctctgtcctgaat

gacattctgagcagactggacaaggtcgaagccgaggtgcagatcgacagactgatcaccggcaggctgcaaagtct

gcagacatacgtgacgcagcagctgattagagccgccgaaatccgggcatctgcaaatctggctgccacaaagatgt

ccgagtgcgtgctggggcagagcaagagagtcgacttctgcggcaaaggctaccacctgatgagcttcccccagtct

gccccgcacggcgtggttttcctgcatgtgacctacgtgcccgctcaggagaaaaatttcaccaccgcccctgccatttg

ccacgacggaaaggcccacttccccagagagggcgttttcgtgagcaacggcacccactggttcgtgacacagagaa

acttctacgagcctcagattatcaccaccgataacacattcgtgtccggcaactgcgacgtggtgatcggcatcgtgaat

aacacagtgtacgaccctctgcagcccgagctggatagcttcaaggaagagctggacaaatacttcaagaaccacacc

agccccgatgtggacctgggcgatatctctggaatcaacgccagcgtggtgaacatccagaaggaaatcgatagactc

aacgaggtggccaaaaacctgaacgagagcctgatcgacctccaagagctgggcaagtacgagcagtatatcaagtg

gccttggtacatctggctgggcttcatcgccggactgatcgctatcgtgatggtgaccatcatgctgtgctgtatgacttct
tgctgcagctgtctgaagggttgttgctcttgcggctcctgctgctga

atgggccaggctgttaccaccccccttaagtttgactttagaccactggaaggatgtcgaacggacagcccac
aacctgtcggtagaggttagaaaaaggcgctgggttacattctgctctgcagaatggccaaccttcaacgtcg
gatggccacgagacggcacttttaacccagacattattacacaggttaagatcaaggtcttctcacctggccc
acatggacatccggatcaggtcccctacatcgtgacctgggaagctatagcagtagaccccccctccctgggt
cagacccttcgtgcaccctaaacctcccctctctcttccccccttcagccccctctctcccacctgaacccccact
ctcgaccccgccccagtcctccctctatccggctctcacttctcctttaaacaccaaacctaggcctcaagtcct
tcctgatagcggaggaccactcattgatctactcacggaggaccctccgccttaccgggacccagggccac
cctctcctgacgggaacggcgatagcggagaagtggcccctacagaaggagcccctgacccttccccaat
ggtatcccgcctgcggggaagaaaagaacccccgtggcggattctactacctctcaggcgttccccttcg
cctgggagggaatggacagtatcaatactggccattttcctcctctgacctctataactggaaaaataacaacc
cctctttctccgaggacccagctaaattgacagctttgatcgagtccgttctccttactcatcagcccacttggg
atgactgccaacagctattagggaccctgctgacgggagaagaaaaacagcgagtgctcctagaggcccg
aaaggcggttcgaggggaggacggacgcccaactcagctgcccaatgacattaatgatgcttttcccttgga
acgtcccgactgggactacaacacccaacgaggtaggaaccacctagtccactatcgccagttgctcctag
cgggtctccaaaacgcgggcagaagccccaccaatttggccaaggtaaaagggataacccagggaccta
atgagtctccctcagcctttttagagagactcaaggaggcctatcgcagatacactccttatgaccctgaggac
ccagggcaagaaaccaatgtggccatgtcattcatctggcagtccgccccggatatcgggcgaaagttaga
gcggttagaagatttgaagagtaagaccttaggagacttagtgagggaagctgaaaagatctttaataaacga
gaaaccccggaagaaagagaggaacgtattaggagagaaacagaggaaaaggaagaacgccgtaggg
cagaggatgtgcagagagagaaggagagggaccgcagaagacatagagaaatgagtaagttgctggcta

ctgtcgttagcgggcagagacaggatagacagggaggagagcgaaggaggccccaactcgaccacgac

cagtgtgcctactgcaaagaaaagggacattgggctagagattgccccaagaagccaagaggaccccgg

ggaccacgaccccaggcctccctcctgaccttagacgattagggaggtcagggtcaggagcccccccctg

**Gag/Pol** aacccaggataaccctcagagtcggggggcaacccgtcaccttcctagtggatactggggcccaacactcc

gtgctgacccaaaatcctggacccctaagtgacaagtctgcctgggtccaaggggctactggagggaagc

ggtatcgctggaccacggatcgccgagtgcacctagccaccggtaaggtcacccattctttcctccatgtacc

agattgcccctatcctctgctaggaagagatttgctgactaaactaaaagcccaaattcactttgagggatcag

gagctcaggttgtgggaccaatgggacagcccctgcaagtgctgaccctaaacatagaagatgagtatcgg

ctacatgagacctcaaaagggccagatgtgcctctagggtccacatggctctctgattttccccaggcctggg

cagaaaccgggggcatggggctggccgttcgccaagctcctctgatcatacctctgaaggcaacctctacc

cccgtgtccataaaacaataccccatgtcacaagaagccagactggggatcaagccccacatacagagact

gctggatcagggaattctggtaccctgccagtcccccttggaacacgcccctgctacccgttaagaaaccgg

ggactaatgattataggcctgtccaggatctgagagaagtcaacaagcgggtggaagacatccaccccacc

gtgcccaacccttacaacctcttgagcgggctcccaccgtcccaccagtggtacactgtgcttgacttaaaag

atgcctttttctgcctgagactccaccccaccagtcagtctctcttcgcctttgagtggagagatccagagatgg

gaatctcaggacaattaacctggaccagactcccgcagggtttcaaaaacagtcccaccctgtttgatgaagc

cctgcacagggacctcgcagacttccggatccagcacccagacctgattctgctccagtatgtagatgactta

ctgctggccgccacttctgagcttgactgtcaacaaggtacgcgggccctgttacaaaccctaggggacctc

ggatatcgggcctcggccaagaaagcccaaatttgccagaaacaggtcaagtatctggggtatcttctaaaa

gagggtcagagatggctgactgaggccagaaaagagactgtgatggggcagcctactccgaagacccctc

gacaactaagggagttcctagggacggcaggcttctgtcgcctctggatccctgggtttgcagaaatggcag

ccccccttgtaccctctcaccaaaacggggactctgtttgagtggggcccagaccagcaaaaggcctaccaa

gagatcaagcaggctctcttaactgcccctgccctgggattgccagacttgactaagcccttcgaactttttgtt gacgagaagcagggctacgccaaaggtgtcctaacgcaaaaactggggccttggcgtcggccggtggcc

tacctgtccaaaaagctagacccagtggcagctgggtggccccttgcctacggatggtagcagccatcgc

cgttctgaccaaagacgctggcaagctcaccatgggacagccactagtcattctggccccccatgcagtaga

ggcactagttaagcaacccctgatcgctggctctccaacgcccgaatgacccactaccaggctctgcttctg

gacacggaccgagtccagttcggaccaatagtggccctaaacccagctacgctgctccctctacctgagga

ggggctgcaacatgactgccttgacatcttggctgaagcccacggaactagaccagatcttacggaccagc

ctctcccagacgctgaccacacctggtacacagatgggagcagcttcctgcaagaggggcagcgcaaggc

cggagcagcagtaaccaccgagaccgaggtagtctgggccaaagcactgccagccgggacatcggccc

aaagagctgagttgatagcgctcacccaagccttaaaaatggcagaaggtaagaagctgaatgtttacaccg

atagccgttatgcttttgccactgcccatattcacggagaaatatatagaaggcgcgggttgctcacatcagaa

ggaaaagaaatcaaaaataaggacgagatcttggccctactgaaggctctcttcctgcccaaaagacttagc

ataattcattgcccgggacatcagaagggaaaccgcgcggaggcaaggggcaacaggatggccgaccaa

gcggcccgagaagtagccactagagaaactccagagacttccacacttctgatagaaaattcagccccctat

actcatgaacattttcactatacggtgactgacataaaagatctgactaaactaggggccacttatgacgatgc

aaagaagtgttgggtttatcagggaaagcctgtaatgcctgatcaattcacctttgaactattagattttcttcatc

aattgacccacctcagtttctcaaaaacaaaggctcttctagaaaggaactactgtccttattacatgctgaacc

gggatcgaacgctcaaagacatcactgagacttgccaagcctgtgcacaggtcaatgccagcaagtctgcc

gtcaaacaagggactagagttcgagggcaccgacccggcacccactgggaaattgatttcactgaggtaaa

acctggcctgtatgggtataaatatcttttagttttcatagacactttctctggatgggtagaagctttcccaacca

agaaagaaactgccaaagttgtaaccaagaagctactagaagaaatcttccccagattcggcatgccacag

gtattgggaaccgacaatgggcctgccttcgtctccaaggtaagtcagacagtagccgatttactgggggttg

attggaaactacattgtgcttacagaccccagagttcaggtcaggtagaaagaatgaataggacaatcaagg

agactttaactaaattgacgcttgcaactggctctagggactgggtgctcctgcttcccctagccctgtatcga

gcccgcaacacgccgggcccccatggtctcaccccatatgaaatcttatatggggcacccccgccccttgta
aacttccctgatcctgacatggcaaaggttactcataacccctctctccaagcccatttacaggcactctacctg
gtccagcacgaagtctggagaccgttggcggcagcttaccaagaacaactggaccggccggtagtgcctc
acccctttccgagtcggtgacacagtgtgggtccgcagacaccaaactaaaaatctagaaccccgctggaaa
ggaccttataccgtcctactgactacccccaccgctctcaaagtggacggcattgcagcgtggatccacgct
gcccacgtaaaggctgccgacaccaggattgagccaccatcggaatcgacatggcgtgttcaacgctctca
aaatcccctaaagataagattgacccgcgggacctcctaa

**Luciferase**

atggaagacgccaaaaacataaagaaaggcccggcgccattctatccgctggaagatggaaccgctggag
agcaactgcataaggctatgaagagatacgccctggttcctggaacaattgcttttacagatgcacatatcga
ggtggacatcacttacgctgagtacttcgaaatgtccgttcggttggcagaagctatgaaacgatatgggctg
aatacaaatcacagaatcgtcgtatgcagtgaaaactctcttcaattctttatgccggtgttgggcgcgttatttat
cggagttgcagttgcgcccgcgaacgacatttataatgaacgtgaattgctcaacagtatgggcatttcgcag
cctaccgtggtgttcgtttccaaaaaggggttgcaaaaaattttgaacgtgcaaaaaaagctcccaatcatcca
aaaaattattatcatggattctaaaacggattaccagggatttcagtcgatgtacacgttcgtcacatctcatcta
cctcccggttttaatgaatacgattttgtgccagagtccttcgatagggacaagacaattgcactgatcatgaac
tcctctggatctactggtctgcctaaaggtgtcgctctgcctcatagaactgcctgcgtgagattctcgcatgcc
agagatcctatttttggcaatcaaatcattccggatactgcgattttaagtgttgttccattccatcacggttttgga
atgtttactacactcggatatttgatatgtggatttcgagtcgtcttaatgtatagatttgaagaagagctgtttctg
aggagccttcaggattacaagattcaaagtgcgctgctggtgccaaccctattctccttcttcgccaaaagcac
tctgattgacaaatacgatttatctaatttacacgaaattgcttctggtggcgctcccctctctaaggaagtcggg
gaagcggttgccaagaggttccatctgccaggtatcaggcaaggatatgggctcactgagactacatcagct
attctgattacacccgaggggggatgataaaccgggcgcggtcggtaaagttgttccatttttttgaagcgaagg

ttgtggatctggataccgggaaaacgctgggcgttaatcaaagaggcgaactgtgtgtgagaggtcctatga
ttatgtccggttatgtaaacaatccggaagcgaccaacgccttgattgacaaggatggatggctacattctgga
gacatagcttactgggacgaagacgaacacttcttcatcgttgaccgcctgaagtctctgattaagtacaaag
gctatcaggtggctcccgctgaattggaatccatcttgctccaacaccccaacatcttcgacgcaggtgtcgc
aggtcttcccgacgatgacgccggtgaacttcccgccgccgttgttgttttggagcacggaaagacgatgac
ggaaaaagagatcgtggattacgtcgccagtcaagtaacaaccgcgaaaaagttgcgcggaggagttgtgt
ttgtggacgaagtaccgaaaggtcttaccggaaaactcgacgcaagaaaaatcagagagatcctcataaag
gccaagaagggcggaaagatcgccgtgtaa

151
